# LayNii: A software suite for layer-fMRI

**DOI:** 10.1101/2020.06.12.148080

**Authors:** Laurentius (Renzo) Huber, Benedikt A. Poser, Peter A. Bandettini, Kabir Arora, Konrad Wagstyl, Shinho Cho, Jozien Goense, Nils Nothnagel, Andrew Tyler Morgan, Job van den Hurk, Anna K Müller, Richard C. Reynolds, Daniel R. Glen, Rainer Goebel, Omer Faruk Gulban

## Abstract

High-resolution fMRI in the sub-millimeter regime allows researchers to resolve brain activity across cortical layers and columns non-invasively. While these high-resolution data make it possible to address novel questions of directional information flow within and across brain circuits, the corresponding data analyses are challenged by MRI artifacts, including image blurring, image distortions, low SNR, and restricted coverage. These challenges often result in insufficient spatial accuracy of conventional analysis pipelines. Here we introduce a new software suite that is specifically designed for layer-specific functional MRI: LayNii. This toolbox is a collection of command-line executable programs written in C/C++ and is distributed open-source and as pre-compiled binaries for Linux, Windows, and macOS. LayNii is designed for layer-fMRI data that suffer from SNR and coverage constraints and thus cannot be straightforwardly analyzed in alternative software packages. Some of the most popular programs of LayNii contain ‘layerification’ and columnarization in the native voxel space of functional data as well as many other layer-fMRI specific analysis tasks: layer-specific smoothing, model-based vein mitigation of GE-BOLD data, quality assessment of artifact dominated sub-millimeter fMRI, as well as analyses of VASO data.

**Highlights:**

- A new software toolbox is introduced for layer-specific functional MRI: LayNii.
- LayNii is a suite of command-line executable C++ programs for Linux, Windows, and macOS.
- LayNii is designed for layer-fMRI data that suffer from SNR and coverage constraints.
- LayNii performs layerification in the native voxel space of functional data.
- LayNii performs layer-smoothing, GE-BOLD deveining, QA, and VASO analysis.

Graphical abstract

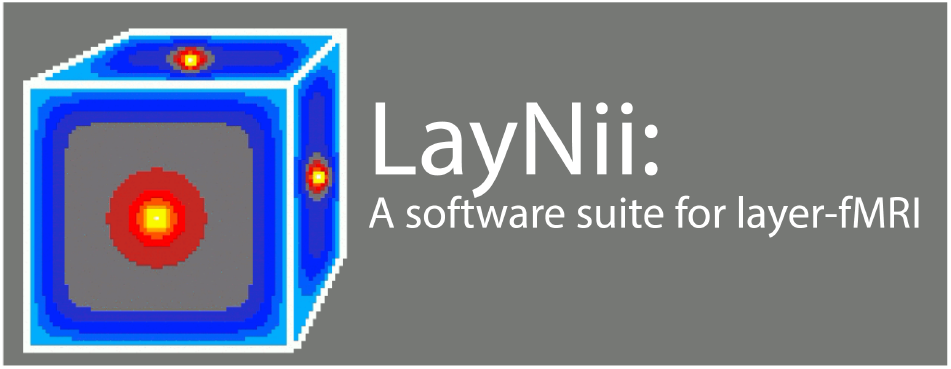

## 1. Introduction

Functional Magnetic Resonance Imaging (fMRI) aims to measure correlates of brain activity at the level of individual voxels. Recent advances in fMRI hardware, readout strategies, and fMRI contrasts provide estimates of brain activation with voxel-sizes in the sub-millimeter domain (Goense et al. 2016). Thus, modern fMRI data from ultra-high fields (UHF) are approaching the spatial scale of cortical layers and columns (Bollmann and Barth 2020; Goense et al. 2012a; Kuehn and Sereno 2018; Petro and Muckli 2017; van der Zwaag et al. 2016). While the first decade of sub-millimeter fMRI research focused on data acquisition methods (Budde et al. 2014; Goense et al. 2010; Goense and Logothetis 2006; Goense et al. 2007; Petridou et al. 2013; Rua et al. 2017; van der Zwaag et al. 2009), the methodological research questions of the layer-fMRI field have since shifted towards addressing analysis challenges. As such, a recent survey of the ISMRM study group *Current Issues in Brain Function* showed that most high-resolution fMRI researchers consider analysis challenges to be more relevant than acquisition challenges (Huber for ISMRM SG CIBF 2018).

The specific shortcomings of common analysis approaches for high-resolution fMRI are listed in multiple review articles (Kemper et al. 2018; Polimeni et al. 2018), and there are many fMRI analysis software packages that are able to minimize these shortcomings to some degree: AFNI/SUMA (https://afni.nimh.nih.gov, (Cox 1996)), ANTs (http://stnava.github.io/ANTs/, (Avants et al. 2008)), BrainVoyager (http://www.brainvoyager.com, (Goebel 2012), CBSTools/Nighres (http://www.nitrc.org/projects/cbs-tools/, (Bazin et al. 2014), https://nighres.readthedocs.io/, (Huntenburg et al. 2018)), FreeSurfer (https://surfer.nmr.mgh.harvard.edu, (Fischl 2012), FSL (http://fsl.fmrib.ox.ac.uk/fsl, (Jenkinson et al. 2012), LIPSIA (https://www.cbs.mpg.de/institute/software/lipsia, (Lohmann et al. 2000)), and SPM (http://www.fil.ion.ucl.ac.uk/spm (Ashburner 2012)). Some of these packages, however, were originally developed, validated and established for the application for fMRI data that have resolutions in the range of 1.5-5mm, large brain coverage and 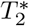-weighted fMRI contrasts. Such fMRI data constitute the vast majority of the fMRI literature and are hereafter referred to as “conventional fMRI”. However, layer-fMRI data are different from conventional fMRI data and layer-fMRI data are limited by many unique high-resolution constraints. Thus, layer-fMRI data cannot be straightforwardly analyzed with conventional analysis pipelines (Fig. 1). The specific layer-fMRI data constraints are listed below:

**Figure 1.**
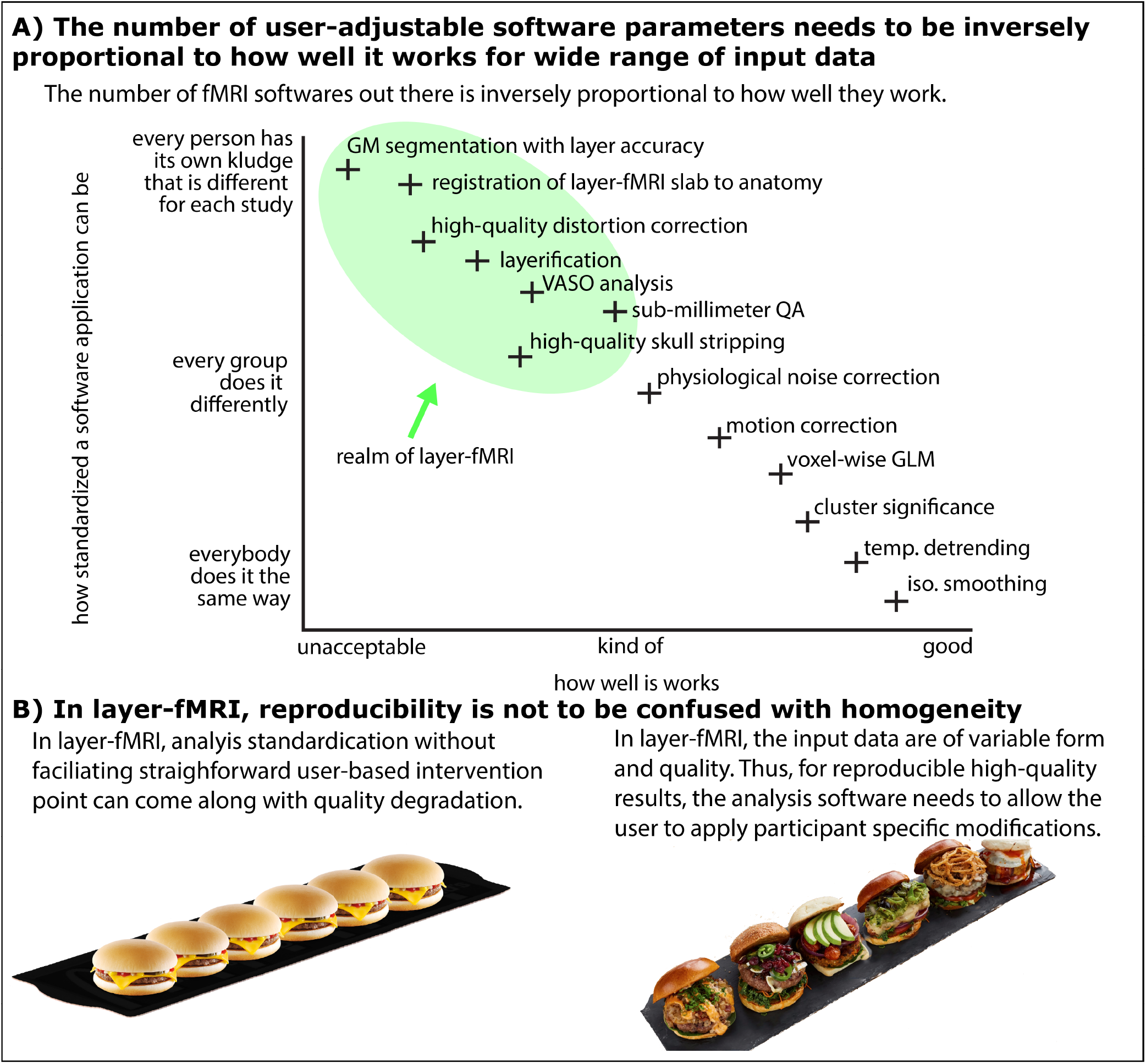
There is a need for a new software suite that is explicitly developed for layer-fMRI. Panel **A)** illustrates that because layer-fMRI data suffer from countless shortcomings, the corresponding processing software needs to give the user the flexibility and responsibility to harmonize analysis algorithm parameters for their specific data’s features. Panel **B)** illustrates that for layer-fMRI data, the possibility and necessity of user dependent algorithm tweaking can result in reproducible results of higher quality as opposed to enforcing standardized homogeneous user-independent black-box analysis pipelines without any user interventions necessary. This case-specific analysis tweaking required broad knowledge of the user about the algorithm and all its tuning parameters.

- **Low SNR:** Layer-fMRI data are noisier compared to conventional fMRI data. In layer-fMRI studies, tSNR values in the single-digit regime are not uncommon. This indicates that layer-fMRI data are usually more limited by thermal noise than physiological noise (Triantafyllou et al. 2011). Therefore, many quality assessment (QA) tools that have originally been developed for conventional fMRI data are not applicable to layer-fMRI data. For example, in the thermal noise dominated regime, the QA metric of tSNR cannot capture many sources of MR-sequence related and physiological-noise-related signal clutter. These sources of signal instability are below the thermal noise floor (for more discussions of inappropriate QA metrics see section 2.5). Thus, for layer-fMRI data, new analysis tools with additional (and optimized) QA metrics are needed. Due to the low tSNR in layer-fMRI, it can become necessary to average multiple data points across space (e.g. spatial smoothing, or voxel pooling). In many layer-fMRI application studies, it can be assumed that neighboring columnar structures are performing the same neural task (e.g. top-down attention) with the identical layer-specific processing signature (albeit potentially different feature representations). In those cases, it can be beneficial to apply local smoothing within the layer direction and without signal blurring across layers (Blazejewska et al. 2019) to enhance the layer-specific processing signature. Alternatively, it can be beneficial to apply intracortical non-isotropic smoothing based on the directional functional activation (Lohmann et al. 2018) or based on the non-isotropic MRI signal intensity across the cortical depth. Thus, for layer-fMRI analyses, new software tools with a larger variety of anatomically informed spatial smoothing methods are needed.
- **Necessity to manually intervene: Accuracy and precision wins over streamlining** In layer-fMRI, the data acquisition procedure is usually pushed to its limits. This means that every acquisition protocol -and image data from each participant-may suffer from individual image artifacts and other experimental short-comings. While minimal manual tuning of conventional analysis pipelines provides more than sufficient quality for conventional fMRI resolutions, it is not uncommon in layer-fMRI applications to invest 8-12 work hours per dataset for manual high-quality corrections of the segmented borders. Furthermore, it is not uncommon that the experimenter needs to spend more than an entire work week per participant to complete an analysis pipeline. This is due to the layer-fMRI specific requirement to have exceptional accuracy of the analysis and high precision of gray matter (GM) borders. This user-dependent layer-fMRI analysis paradigm is in direct opposition to the demands of large scale population studies of conventional fMRI (UKBiobank, HCP, etc.). In those large scale studies, the analysis pipelines need to be executable with minimal user interventions; and in those population studies, analysis robustness can outrank small accuracy tradeoffs. While these standardized conventional fMRI analyses have gone through several decades of protocol optimizations and aims for stable and robust streamlined analysis that can process thousands of participants without necessary user-interventions, this is neither possible, nor currently demanded for layer-fMRI analyses. In layer-fMRI, the researcher needs to have the flexibility to specifically adjust the analysis pipeline for each individual data set. Thus, a timely layer-fMRI analysis toolbox should be modular and optimized to allow straightforward user-based manipulations of a wide range of algorithm parameters. This would aid the user to iteratively tweak the parameters for every dataset to end up with fine-tuned pipelines that provide the highest possible quality. Such an approach is necessary to optimize layer-fMRI analysis strategies globally in the long term until the data quality will allow streamlined analyses.
- **Restricted coverage:** Layer-fMRI acquisition protocols are usually pushed to the limits of what the scanner can achieve. Thus, high spatial resolutions are often achieved by significant tradeoffs in coverage and even single-slice protocols are accepted as a compromise for resolution (Cheng et al. 2001; Yacoub et al. 2008). These coverage constraints in high resolution fMRI are usually due to the limitations of the MR hardware (mainly gradient performance) and the need for a TR in the range of seconds to allow statistical analysis of the fMRI signal. Furthermore, layer-fMRI data are often acquired with small receiver arrays that have limited coverage and do not facilitate straightforward whole-slice analyses. While it is without question that small fMRI coverage can be troublesome for many analyses, small coverage data play a crucial part in this rapidly developing field. However, many conventional fMRI analysis pipelines are not designed for (or are incompatible with) small coverage data. Therefore, new layer-fMRI analysis tools that are more accommodating to small coverage data are needed.
- **No topology requirements of the cortical sheet:** Almost all current layer-fMRI studies use reduced field of views (Schluppeck et al. 2018) and do not exceed a coverage volume beyond a thin slab. However, many of the current topography-related analysis packages have been designed for whole brain analyses that need to fulfill a number of topological requirements. Namely, fMRI data are used in brain models of closed, continuous GM sheets, without holes or crossing surfaces. However, even though the biological structure of the cerebral cortex fulfils this requirement, common layer-fMRI EPI slab-data do not fulfill this requirement. It is not uncommon for some parts of the functional data to suffer from holes in the cortical sheet and not fulfill the original topology requirements. There are discontinuities, resolution constraints, as well as constraints of brain coverage (commonly 0.5-2.5 cm slabs) that violate the topology constraints in virtually all layer-fMRI data. Therefore, there is a need for analysis software that allows topographical analysis of layers and columns with fewer topological requirements (Kemper et al. 2018).
- **Necessity to work in the data type of voxels:** There are various philosophical approaches to layer-fMRI analyses. Many brain researchers prefer to analyze, depict and interpret their data in a data format that best resembles the object of interest. Since the brain consists of a folded sheet of stacked layers, one could analyze and depict brain data in surface space (using mesh vertices). Alternatively, many MRI and reconstruction methodologists prefer to keep their data as raw (and original) as possible and thus prefer to analyze, depict and interpret their data in the format they are acquired (in voxel space) and without additional necessary data conversions. Since layer-fMRI is still largely limited by the data acquisition strategies, it might be appropriate to favor data processing tools that operate in voxel space to avoid additional confounds that come from transitioning to surface space. Thus, in layer-fMRI, there is a need for purely voxel-based analysis software in addition to already existing surface-based analyses. If researchers had access to layering analyses in voxel space, the data acquisition and analysis could be brought closer together. This would allow researchers to optimize acquisition and analysis concurrently and would enable a new form of fMRI studies that were still unthinkable until today. For example, with layer estimates directly accessible in the raw scanner voxel space, online analysis at the scanner console are doable and layer-dependent neural-feedback studies can become possible.
- **No anatomical reference requirements:** Since layer-fMRI data are limited by a high noise level, a successful study often depends on extensive averaging of countless task trials. In light of limited research funding for expensive scan time and ethically appropriate finite scan durations, it is often challenging for the experimenter to acquire additional non-fMRI auxiliary data that might aid the analysis. As such, it can be challenging to obtain high-quality high-resolution whole brain anatomical reference data, *B*_0_ and *B*_1_ maps, as well as reliable distortion inverted EPI reference data. It is also not uncommon that a researcher is confronted with the decision: either a) to obtain many auxiliary analysis-facilitating reference data without remaining scan time for sufficient functional averages, or b) to obtain sufficient functional averages for decent functional interpretability but without additional reference data. Furthermore, the authors believe that even in cases when these auxiliary data are available, their utilization at submillimeter accuracy level is rarely perfect. While the lack of reference data is not ideal, there is a need for software analyses that can still extract useful information from layer-fMRI without reference data. Thus, there is a need for a layer-fMRI analysis software that is able to perform layer-specific analyses directly in the distorted native EPI space (Chai et al. 2019; van der Zwaag et al. 2018a;b) without the requirement of additional non-fMRI data.
- **Non-BOLD fMRI contrasts:** At high spatial resolutions, conventional GE-BOLD fMRI sequences are limited by spatially unspecific draining veins that obscure the underlying layer-specific neural activation. Thus, it is getting popular to use alternative fMRI contrasts that do not suffer this shortcoming (Chai et al. 2019; Huber et al. 2019b). One example is the blood volume sensitive VASO (vascular-space-occupancy) contrast that is used in approximately 30% of current layer-fMRI studies (Huber et al. 2020a). The VASO sequence, however, does not directly provide a single contrast time course like conventional fMRI GE-BOLD data do. Instead, raw VASO time series data consist of multiple interleaved contrasts with variable repetition times (TR) and require additional preprocessing steps to extract pure BOLD and pure blood-volume weighted time courses, respectively. Thus, there is a need for an analysis software that performs respective VASO processing analyses including temporal image resorting, nifti header manipulations and dynamic contrast divisions.
- **Draining vein problem:** In cases where layer-fMRI data have already been acquired with GE-BOLD, the unwanted venous signal can no longer be accounted for with advanced acquisition methods. Instead, one could model the directional blood flow in the cortical vasculature to estimate the venous signal contamination (Markuerkiaga et al. 2016) and remove it (Markuerkiaga et al. 2020) as much as possible. Thus, a timely layer-fMRI analysis software should contain corresponding tools for model-based venous signal removal.

In short, layer-fMRI data often suffer from multiple short-comings that hinder a straightforward analysis of layers and columns. While the acquisition methodology needs to be further advanced to remove those shortcomings, there is a current need for a layer-analysis software that works despite each and all of the current shortcomings (Fig. 1).

Here, we present an fMRI analysis software suite LayNii that is specifically designed for layer-fMRI data and that addresses all of the above listed challenges. LayNii is designed to perform layer analyses entirely in voxel space and consists of many modular programs that perform layer-fMRI specific tasks:

a. Cortical depth and thickness measurements with corresponding assignments of layer tags to each GM voxel (a.k.a. layerification) in any segmented volume data set,
b. Estimation of columnar structures across voxels in any volume data set,
c. Performing various ways of layer-specific smoothing,
d. Providing a set of appropriate QA metrics for thermal noise-limited layer-fMRI data,
e. Pre-processing blood-volume based VASO fMRI contrasts that are optimized for the layer-specific microvascular responses,
f. Applying model-based deveining algorithms of layer-fMRI GE-BOLD data.

## 2. LayNii algorithms

### 2.1. Package structure

The LayNii software suite is available as stand-alone C++ source code without external dependencies or needing libraries. It is distributed as pre-compiled (binary) installation packages for all major operating systems (Linux, Windows, macOS). LayNii can read and write nifti files ( Neuroimaging I nformatics Technology; https://nifti.nimh.nih.gov/) in uncompressed NII and compressed NII.GZ formats.

The name LayNii is derived from the two words “**Lay** erfMRI” and “**Ni** ft**i**”. The prefix *LAY* stresses that this suite is particularly built for layer-fMRI (which includes sub-millimeter fMRI and columnar fMRI). The suffix *NII* emphasizes that this software suite operates in the voxel space of nifti data.

The structure of LayNii is largely inspired by the philosophy of alternative software packages and is aimed to be as modular as possible. It is designed to consist of multiple individual lightweight programs that are executable from the command line and can be combined in pipeline scripts that may or may not include additional elements of alternative software suites like, AFNI, fslmaths, ANTs, etc. LayNii’s modularity will also help to straightforwardly include LayNii’s programs into already established software suites in the future. We follow the AFNI principle of providing mechanisms, not policies. We aim to give the user the power to assemble computing pieces in different ways to make customized analysis, which in turn means that it is the user’s responsibility to know what the individual programs with all its tuning parameters do. This might give LayNii a high flexibility to increase the precision and accuracy for each given data set at the cost of straightforward streamlining. Adjusting LayNii’s tuning parameters can have a large influence on how the final results look. For the sake of reproducibility, we encourage users of LayNii to report the version number (each LayNii release has a unique DOI using Zenodo) and all specifics of the executed LayNii commands (input parameters) in their respective publications. Currently, the aim of the LayNii package is not to provide a one-shot analysis solution for entire pipelines, as this is already well-covered by other available packages. There is version control of LayNii, so researchers can reproduce results from older studies even after several updates.

Some of the most essential LayNii programs are discussed below. An exhaustive list of additional programs and tutorials of all (>30) programs is available on Github. Each individual program has a *-help* option that describes the main functionality and how to use it.

### 2.2. Layering: algorithm description and examples

Layer-fMRI necessitates determining the cortical depth of each gray matter voxel. This information can be used to pool together every voxel within a certain cortical depth range to generate “layers”. Since the cortex varies in thickness across regions, absolute (as opposed to relative) cortical depth measurements are often not very useful to consistently determine layers. To account for this variation across regions, local cortical thickness measurements are used to normalize the cortical depths. We refer to the resulting normalized cortical depth measurement as “equi-distant metric”. The equi-distant metric yields equi-distant layers, when quantized.

In addition to cortical thickness, the location and size of the cortical neurobiological layers also vary with regards to the cortical curvature. The effect of cortical curvature on layering was originally demonstrated by Bok in 1929, where he subdivided the cortex into columns to show that deeper layers get thicker and superficial layers g et thinner around gyri. Meanwhile, deeper layers get thinner and superficial layers get thicker around s ulci. This observation is known as the equi-volume principle (Waehnert et al. 2014).

In LayNii, we provide both equi-distant and equi-volume layering options to the user. While solutions already exist (Bazin et al. 2014; Fischl 2012; Goebel 2012; Huntenburg et al. 2018; van Mourik et al. 2019; Wagstyl et al. 2018), our implementation differs in a unique way. It is computed completely on the discrete lattice (as opposed to by means of surface approximations) (Glen et al. 2018; Roden 2019). To give a description of our layering algorithm, first we define the terms that will be referred to later (also see Fig. 2)):

**Figure 2.**
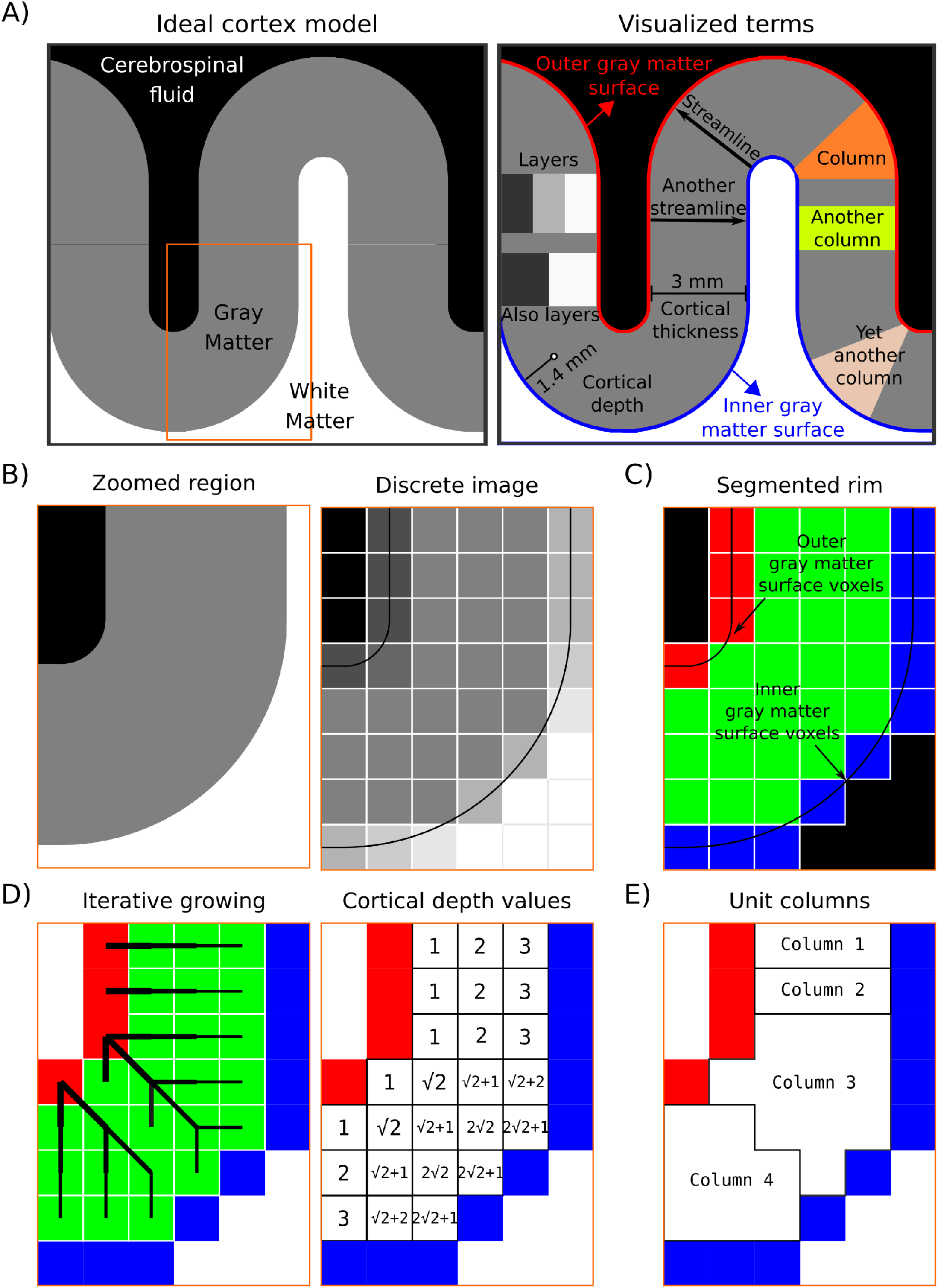
Description of the layerification algorithm. Panel **A)** shows an ideal cortex model consisting of sharp tissue boundaries together with terms exemplified on this model. Panel **B)** shows zoomed-in regions of the model together with a discretized version. This discretized image is analogous to CSF nulled *T*1 - weighted MRI data. Panel **C)** shows the segmentation required by LayNiis layering algorithm. Black voxels indicate the irrelevant voxels, blue voxels indicate the inner gray matter surface, red voxels indicate the outer gray matter surface voxels, and green voxels indicate the pure gray matter voxels. Panel **D)** visualizes the spatial intuition behind measuring the cortical depth together with showing the measured cortical depths. Note that here the cortical depths are only measured from the outer gray matter surface voxels (red), however LayNii also measures the cortical depth relative to the inner gray matter surface voxels (blue). Therefore, each pure gray matter voxel (green) is described by two cortical depths. Also note that when added, these two cortical depths measure the cortical thickness. Panel **E)** shows the unit columns defined by LayNii’s algorithm. These unit columns are used to compute the equi-volume metric.

- **Layer(s):** In neuroscience, the term “cortical layers” often refers to neurobiological layers (e.g. histologically defined). Here we are not using layers to exclusively mean neurobiological layers, but instead to mean “a thickness of some voxels laid within cortical gray matter” which may or may not correspond to neurobiological layers. Due to the thickness of neurobiological layers and the typical spatial resolution used in depth dependent fMRI, an actual matching of artificial and neurobiological layers can only be achieved in case of layerification with higher resolutions than the native spatial resolution of the fMRI sequence used.
- **Inner gray matter surface:** Portion of gray matter that mostly faces white matter.
- **Outer gray matter surface:** Portion of gray matter that mostly faces cerebrospinal fluid.
- **Streamline:** A line that connects inner and outer gray matter surfaces based on some principle (not necessarily the shortest Euclidean distance).
- **Cortical thickness:** Shortest streamline distance between outer and inner gray matter borders.
- **Cortical depth:** Distance from inner **or** outer gray matter surface, a portion of cortical thickness. Slightly different from *layer* because this term indicates a **quantitative measure** which can be used to define the neurobiological layers.
- **Column:** A group of **voxels** that are successively penetrated by one or multiple streamlines. Not to be confused with neurobiological cortical columns which indicates a group of neurons, not voxels.
- **Unit column:** Smallest units of columns that can be defined in a discrete lattice (ordered set of voxels) given the image resolution and topology of the cortex. These unit columns are solely used for the purpose of facilitating subsequent layerification. Their definition is chosen to be algorithmically convenient and not intended to be physiologically plausible in all instances. Their width is chosen to be as thin as possible while touching at least one outer gray matter voxel and at least one inner gray matter voxel. This definition of unit columns should not be confused with the definition of cortical distances in section 2.3 (below).
- **Curvature:** A measure of ‘how much a shape bends’. More specifically, here we are defining it as a measure of ‘how much a unit column bends’. E.g. banana-shaped or pyramidal-shaped.
- **Rim:** A volume of cortical gray matter. Here, we specifically use this term to indicate our input image, which consists of four integer values (0=irrelevant voxels, 1=inner gray matter surface voxels, 2=outer gray matter surface voxels, 3=pure gray matter voxels; see panels of Fig. 2).

In what follows, we detail the implementation steps of layerification in LayNii.

1. The layerification algorithm in LayNii starts from a segmented rim image (see Fig. 2) Panel C). This segmented rim image contains the segmentation of GM and its borders. The segmentation itself is not done in LayNii (see section 3.1). The LayNii program `LN_RIMIFY` can convert the segmentation output of alternative software packages to a LayNii-optimized rim file.
2. For each voxel, we measure two cortical depths. One relative to the inner gray matter surface voxel and another relative to the outer gray matter surface voxels. We compute these cortical depths by using an iterative growing algorithm.
3. By adding relative distances to inner and outer gray matter surfaces, we compute the cortical thickness per voxel.
4. Cortical depth relative to inner gray matter surface voxels divided by thickness provides an equi-distant metric (ranges between 0 to 1).
5. One streamline per gray matter voxel (green) is defined by the closest outer (red) and the inner (blue) gray matter surface voxels. We use the streamlines to describe the curvature type (as either gyrus, sulcus or straight wall). This is done by means of counting the unique inner and outer gray matter surface voxels each streamline of a unit column connects to. Note that this method avoids traditional curvature computations while still yielding the useful curvature type information.
6. Then we define our unit columns. A unit column has to touch at least one outer gray matter voxel and at least one inner gray matter voxel. A unit column might correspond exactly to one streamline where the cortex curvature is mostly straight (and thin). However, unit columns will often correspond to multiple streamlines when the cortex is curved (and thick). Such unit columns exhaustively cover all gray matter voxels (see illustrative examples in Fig. 2E). The width of the unit column is as small as the voxel resolution allows. They are defined to have at least one voxel of the inner and at least one voxel of the outer gray matter.
7. By exploiting the equi-distant metric, we count the voxels that fall close to the superficial side versus the deeper side of the middle gray matter surface.
8. In order to ultimately estimate equi-volume layers, we compute how much and to which direction the middle gray matter should be pushed to in order to balance the number of voxels on each side. We call the resulting numbers *equi-volume factors*.
9. Then we exploit the equi-distant metric as a constrained vector space (adding up to always one when the normalized cortical depth relative to inner and outer gray matter surfaces are added together). This constrained vector space can be recognized as a simplex space of two dimensions. The n-dimensional simplex space was first described by August Ferdinand Möbius in 1827 under the name “barycentric coordinates” and brought back to attention in the modern era by John Aitchison in 1986 (Aitchison 1986) under the name *compositional data*. We perform a mathematical translation operation that is defined as a linear operator within the simplex space (called perturbation in Pawlowsky-Glahn et al. (2015)) by using the equi-volume factors. This translation balances the metric space in a way to yield the equi-volume metric. Note that this operation is done on a per unit-column basis.
10. Since the unit-columns have sharp transitions in between, it can be advantageous to smooth these transitions to have a more biologically plausible equi-volume metric distribution over gray matter voxels. We use iterative smoothing with a small kernel to prevent leakage around kissing gyri and tight sulci.
11. As the final step, we quantize the equi-distant and equi-volume metrics to provide the desired number of layers to the user. However, note that we are also providing the equi-distant and equi-volume metrics as default outputs. This is advantageous to allow the users to choose their own quantization methods.

This layering procedure works in isotropic and asymmetric voxels without directionality biases.

Layering in LayNii works directly on 2D and 3D images. In addition, it works equally well for partial coverage and whole brain images. As an example, see top row of Fig. 3 for layering applied on synthetic/simulated a 2D image. For real-live examples of 2D-layerification on individual histological slices and pictures thereof see https://layerfmri.com/2dlayers/. The middle row of Fig. 3 depicts an example of a 3D partial coverage image (cut from respective whole brain data) and bottom row depicts a whole brain image.

**Figure 3.**
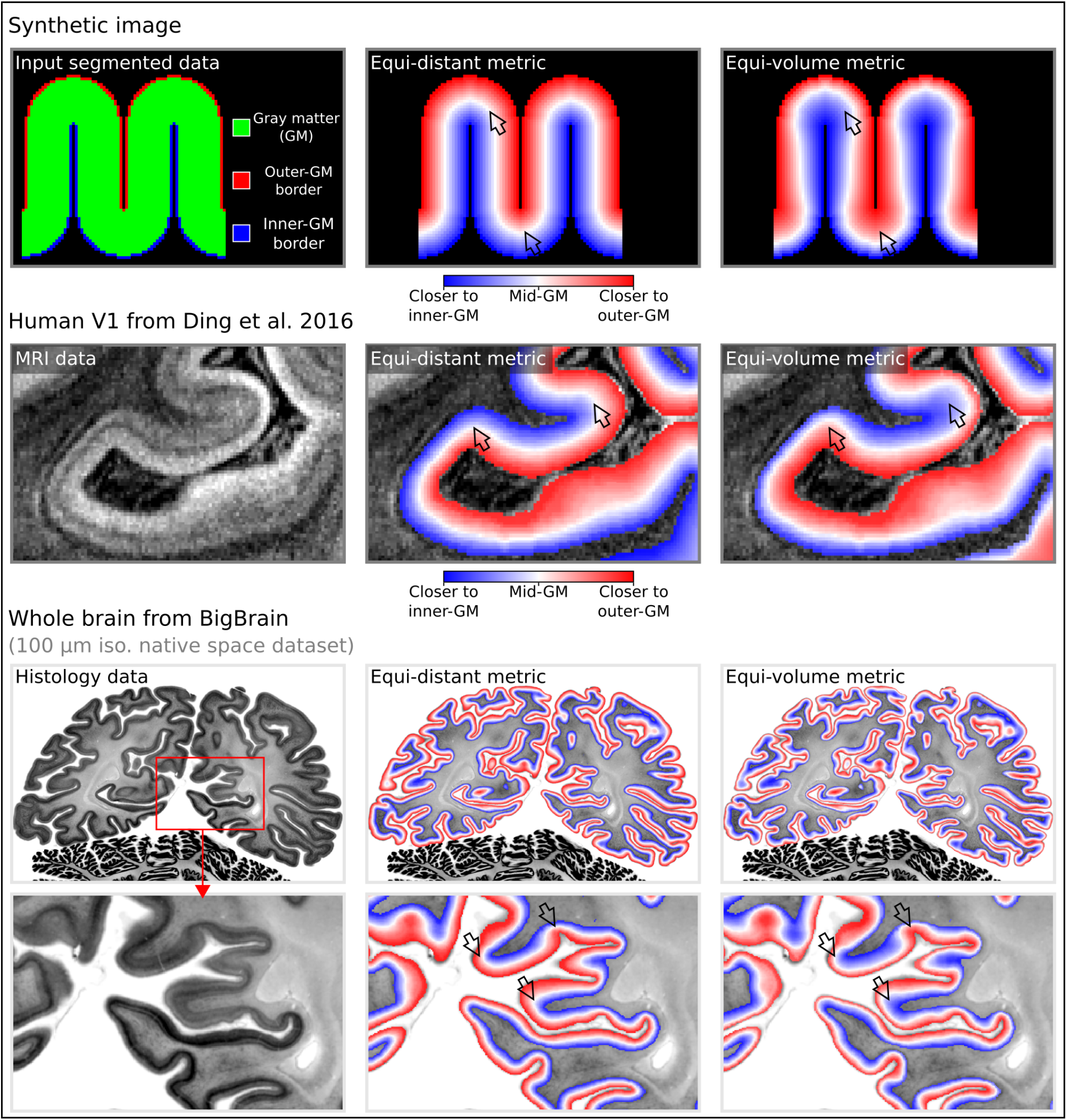
Layering metrics generated in LayNii. The top row shows an application with a synthetic 2D image. The middle row shows the empirical layers from Ding et al. (2016) (0.2 mm iso.). The bottom row shows BigBrain (0.1 mm iso., native space) (Amunts et al. 2013) with cortical borders provided in Wagstyl et al. (2020). The equi-distant metric is shown in the middle column and equi-volume metric is shown in the right column for each image type. To better appreciate the difference between the equi-volume and equi-distance layers on the BigBrain data, see the gif animation in Fig. 6) online: https://thingsonthings.org/ln2_layers/. The arrows highlight areas where the equi-distant and the equi-volume metric differ considerably.

Since the LayNii algorithms are operating directly in voxel space, the layerification can be significantly faster than conventional approaches. For instance, the duration for whole-brain 0.5 mm iso. data processing on a single CPU takes approximately 8 sec for equi-distancing and 48 sec for equi-voluming. The corresponding computation times for 0.2 mm whole brain data (BigBrain) takes 2 and 12 min respectively. For more information on the computation times, see: https://thingsonthings.org/ln2_layers/. For a practical (video-) tutorial of performing layerification directly on layer-fMRI EPI data see here: https://layerfmri.com/analysispipeline/ and here: https://layerfmri.com/quick-layering/. For an in-depth discussion and comparison of equi-distance layering, equi-volume layering and additional layering algorithms that are implemented in LayNii see: https://layerfmri.com/equivol/.

Fig. 4 depicts examples of the layerification in native distorted EPI space. While it is often advantageous to perform the layerification directly in the native distorted E PI space, it is not necessary (nor advised) to perform the layerification in the same spatial grid size. Depending on the number of extracted layers, it can be advantageous to perform the layerification of a finer spatial voxel grid. This can be done, while leaving the raw fMRI data in the original spatial grid and is comparable to vertex coordinates that are defined with finder precision as the voxels. For more hand on examples of how to do this with LayNii, see https://layerfmri.com/how-many-layers-should-i-reconstruct/.

**Figure 4.**
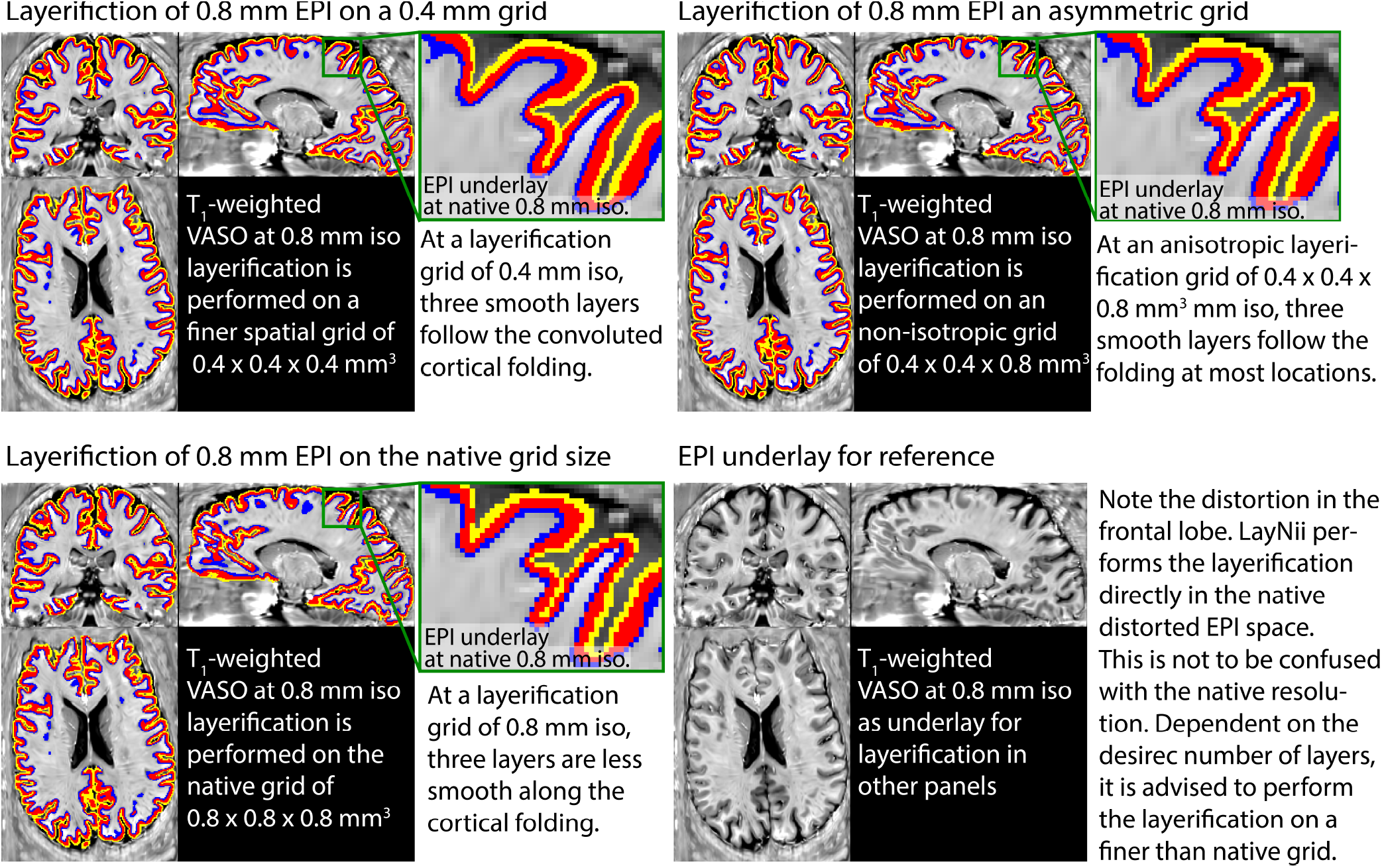
Examples of the layerification in native distorted EPI space. Example of performing layerification on the whole brain fMRI at 0.8 mm isotropic resolutions. Here, an example is shown with three extracted layers. Typically, at 0.8mm resolutions, not more than 2-3 layers can be extracted without losing smoothness along the three dimensional cortical folding (bottom left panel). However, when the layerification is performed on a finer spatial grid, the smoothness is improved (top panels).

### 2.3. Estimation of columnar distances

While voxels with the same relative distance to the GM surfaces are collected into bins of layers, it can also be possible to group voxels into bins of the same columnar distances, in the orthogonal direction to layers, where the distances are measured from specified landmarks. Such estimates of columnar structures that span across all layers of the cortical depth are also often needed for many steps of layer-fMRI analysis. That is, in studies where layer-specific activity cannot be assumed to be identical across large patches of GM, it becomes necessary to perform simultaneous laminar and columnar analyses (Goense et al. 2012b; Huber et al. 2020b; Moerel et al. 2018b; 2015; Olman et al. 2012). As such, estimates of columnar distances are vital for research questions addressing the topographical distributions of layer-dependent brain activation. Past examples are a) depictions of somatotopically aligned body-part representations in the primary sensory cortex (Yu et al. 2019), b) layer-dependent signal distribution across visual eccentricity (Huber et al. 2019a)(p 25), c) for columnar-specific functional hierarchy mapping (Huber et al. 2020a), d) for cortical unfolding (Persichetti et al. 2020), or e) the analysis of tapping induced activation patches that do not extend across a cortical patch of few millimeters in the primary motor cortex (Huber et al. 2020b). In LayNii, columnar distances are calculated in a six-step algorithm that is schematically illustrated in Fig. 5A:

**Figure 5.**
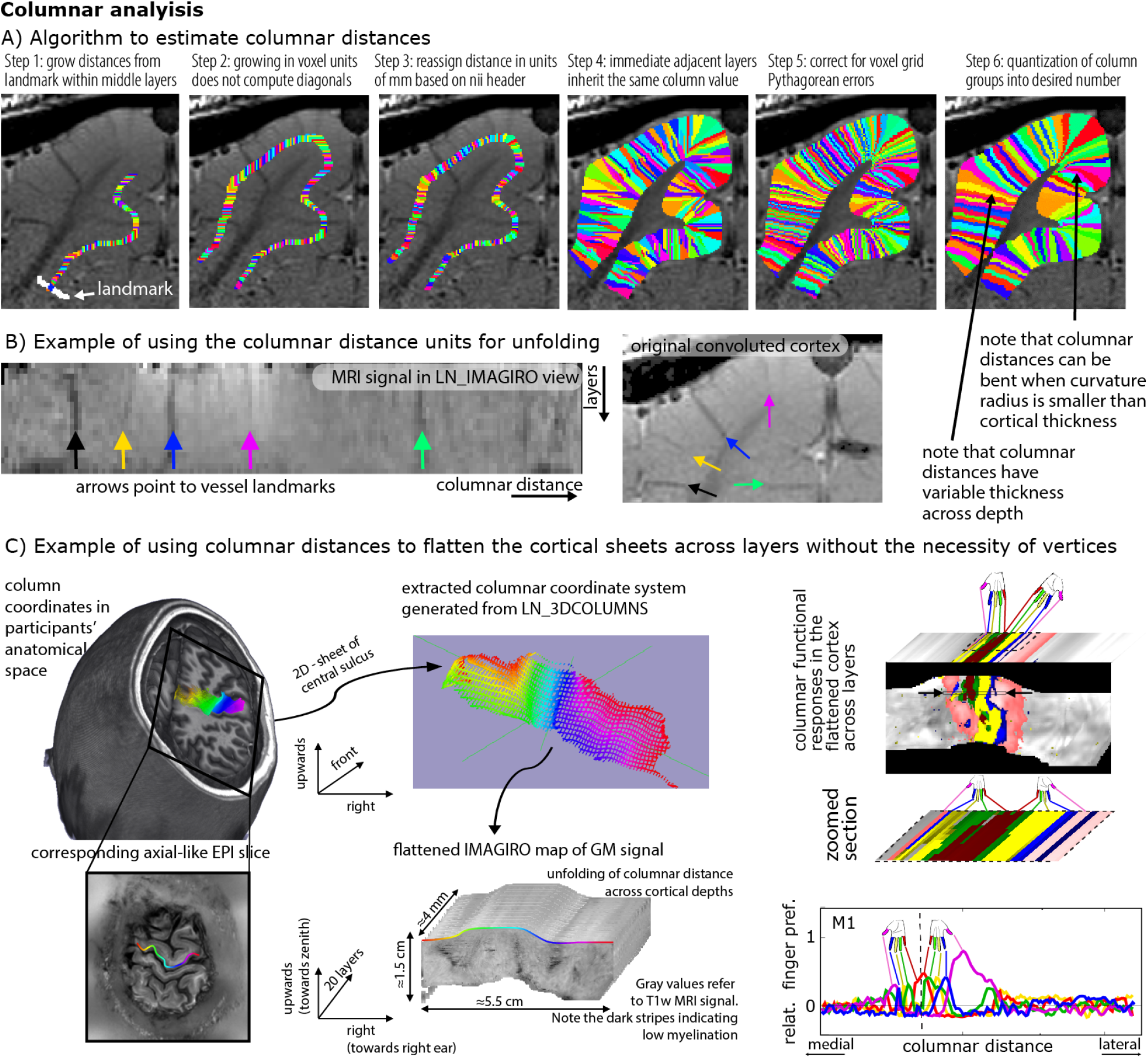
Estimating columnar distances in voxel space with LayNii. Panel **A)** schematically describes the underlying algorithm of LayNii’s columnar distance estimation. Panel **B)** depicts the corresponding MRI signal in two independent coordinate systems: a) the scanner coordinate system with folded GM and b) the unfolded cortical ribbon with orthogonalized depth and column distances. The data presented in panels A-B are acquired with an 8 weeks old female cat, Varian 9.4T at CMRR, resolution: 0.125 × 0.125 × 0.5 mm3, Gradient Echo MultiSlice imaging sequence (GEMS, Agilent technology, Inc.) sequence. Panel **C)** depicts a potential application study of the columnar coordinate system for topographic mapping of functional movement representations. The data presented in panel C are acquired with VASO at a SIEMENS magnetom 7T at FMRIF/NIH with 0.8 mm3 resolution and have been previously described in (Huber et al. 2020b).

1. In contrast to layer estimates, there is usually no clear physiologically defined coordinate system origin. While layer distances are inherently normalized to a coordinate system between WM and CSF borders, the origin of columnar estimates is highly dependent on the specific study and research question. Thus in LayNii, columnar distance estimates are generated based on a manually set landmark (Fig. 5A). This is the starting point of the algorithm (white arrow). In past applications of this algorithm, LayNii users placed the landmarks at anatomical reference points (e.g. position of smallest curvature radius in hand knob) or functional reference points (e.g. position of V1-V2 border, border of thumb finger representation in S1).
2. This landmark is used as the origin of a subsequence growing algorithm. In order to account for the fact that the GM of a neighboring sulcus can be closer than the distance of a column within a sulcus of kissing gyri, the distance is estimated here with a local grow-algorithm that extends the local patch of columnar distances iteratively voxel-by-voxel (step-size=1 voxel).
3. Since the growing algorithm works in voxel space, it can only estimate distances in units of integer multiples of voxel distances. This means that a diagonal voxel neighbour is estimated as being two steps away (one step in each orthogonal direction), whereas the Euclidean distance is actually smaller (square root of two). The corresponding Pythagorean errors are mitigated by the application of an additional processing step that applies local smoothing along the layer direction only.
4. In the next step, the columnar distances are extrapolated across layers. The columnar distance value of every voxel is simply inherited from the next closest voxel that has a determined distance estimate already. This is done within the GM ribbon only (to avoid leakage from neighboring sulci).
5. Again, this step comes along with Pythagorean errors (mismatch of growing iteration steps and Euclidean distance) that can be accounted for with within-layer smoothing.
6. Finally, the user can choose the desired column thickness. This is done with the optional `-Ncolumns` flag of the LayNii program. When this parameter is not specified by the user, the maximum number of columns is used, and the resulting columnar estimates are one voxel thick. For smaller values of `-Ncolumns`, the columns become thicker. The width of the columnar distances is defined to be equal in the middle layer along the cortical ribbon. This means that at locations of strong cortical curvature, the columnar distances have a shape of a frustum of a pyramid exhibiting different widths across the cortical depth.

Note that the columnar distances can be bent in areas of strong cortical folding (Fig. 5A, step 6). While there are cytoarchitectonic (Bok 1929) and angioarchitectonic (Pfeifer 1940) indications that this might be physiologically plausible, a general and quantitative parametrization of the columns bending and shape has not been described in the literature so far. Thus, further research is needed to investigate the physiological accuracy of the columnar bending in LayNii. A more detailed discussion of the columnar distance shape in LayNii is given in: https://layerfmri.com/equivol/.

These columnar distance estimates can then be further used for cortical unfolding Fig. 5)B.

Fig. 5C depicts an example application of LayNii’s columnar distance estimation and subsequent cortical flattening in three dimensional voxel space. Here the mirrored finger representations (Huber et al. 2020b) in the primary motor system are used as a toy-model to exemplify the purpose of the columnar distance analyses in LayNii. The left panels depict the estimates of the columnar coordinates in the native volume space of T1-weighted functional EPI. Here, two orthogonal axes of columnar distances are used. One axis goes along the medial-lateral direction (colors left-right), and one axis goes across the direction of the depth of the central sulcus (upwards-downwards). This spans a locally confined orthogonal grid of columnar distances across the entire anterior bank of the central sulcus and can be used to extract functional signals across all layers (third dimension). These three units of distances (medial-lateral, upwards-downwards, and layers). Can be used to re-grid the signal into a new nifti file with the LayNii program `LN_IMAGIRO`, by simple signal value copying of each voxel to another nifti file based on the three new coordinates. The result is shown on the right panel of Fig. 5C as flattened cortical patches of functional finger dominance maps in 3D-nifti pace. It is shown how the flattened volume representation of the data facilitates straightforward analysis across columnar profiles projected across cortical depths.

A more detailed explanation of the columnarization in the LayNii program `LN_3DCOLUMNS`, and the unfolding program `LN_IMAGIRO` is given here: https://layerfmri.com/columns/.

### 2.4. Layer-specific smoothing

Due to the low tSNR in layer-fMRI, it can become advantageous to apply spatial smoothing. In order not to compromise the locally specific layer-fMRI signal of interest, the spatial smoothing kernel needs to be exclusively applied in anatomically informed directions only. Spatial smoothing in specific layer directions has been originally proposed for vertex-based analyses in (Polimeni et al. 2015), re-implemented for voxel-based layer smoothing in (Huber et al. 2017; 2018), and it has been ultimately described in full depth later (Blazejewska et al. 2019). This form of layer-specific smoothing is commonly applied locally in a way that the signal leakage drops off based on a Gaussian kernel of a given full-width-half-maximum (FWHM), analogous to conventional isotropic smoothing. In contrast to conventional isotropic smoothing, however, the layer-specific smoothing has an additional signal leakage drop off penalty perpendicularly to the layer-direction. Thus, for layer-specific smoothing, the signal leakage is restricted across voxels in different layers, despite the case that they might be very close to each other in Euclidean space. In the LayNii program, `LN_LAYER_SMOOTH` this is implemented as volume smoothing in voxel space by restricting the smoothing kernel across voxels whose centroids are located in the same cortical depth. The smoothing is thus applied within masks of previously defined layers only.

This form of layer-specific smoothing, can be seen as isotropic smoothing within a collection of binary masks (comparable to AFNIs `3dBlurInMask`). So if the user chooses to have very few (but thick) layers, smoothing would be isotropic within each of those layers, with hard boundaries in between layers where no smoothing occurs. On the other hand, when the user would choose to have as many as 100 (very thin) layers, the smoothing algorithm solely smoothes across voxels that are in the same percent-bin of cortical depth. Using more layers, reduces signal leakage across cortical depth within the same depth-bin.

When the smoothing kernel sizes are particularly large and approach the spatial scale of the distance between touching gyri, it becomes necessary for the algorithm to consider signal leakage across sulci in very close proximity (Fig. 6A). While the problem of signal leakage across neighboring gyri is inherently avoided in surface-based smoothing approaches (Jo et al. 2007; Kiebel et al. 2000), it needs to be explicitly taken care of in layer-specific volume smoothing programs of LayNii. In the LayNii implementation of layer-specific smoothing, a specific `-NoKissing` flag ensures that smoothing does not occur blindly across all voxels in close proximity, but only in those voxels that are connected to the same GM area.

**Figure 6.**
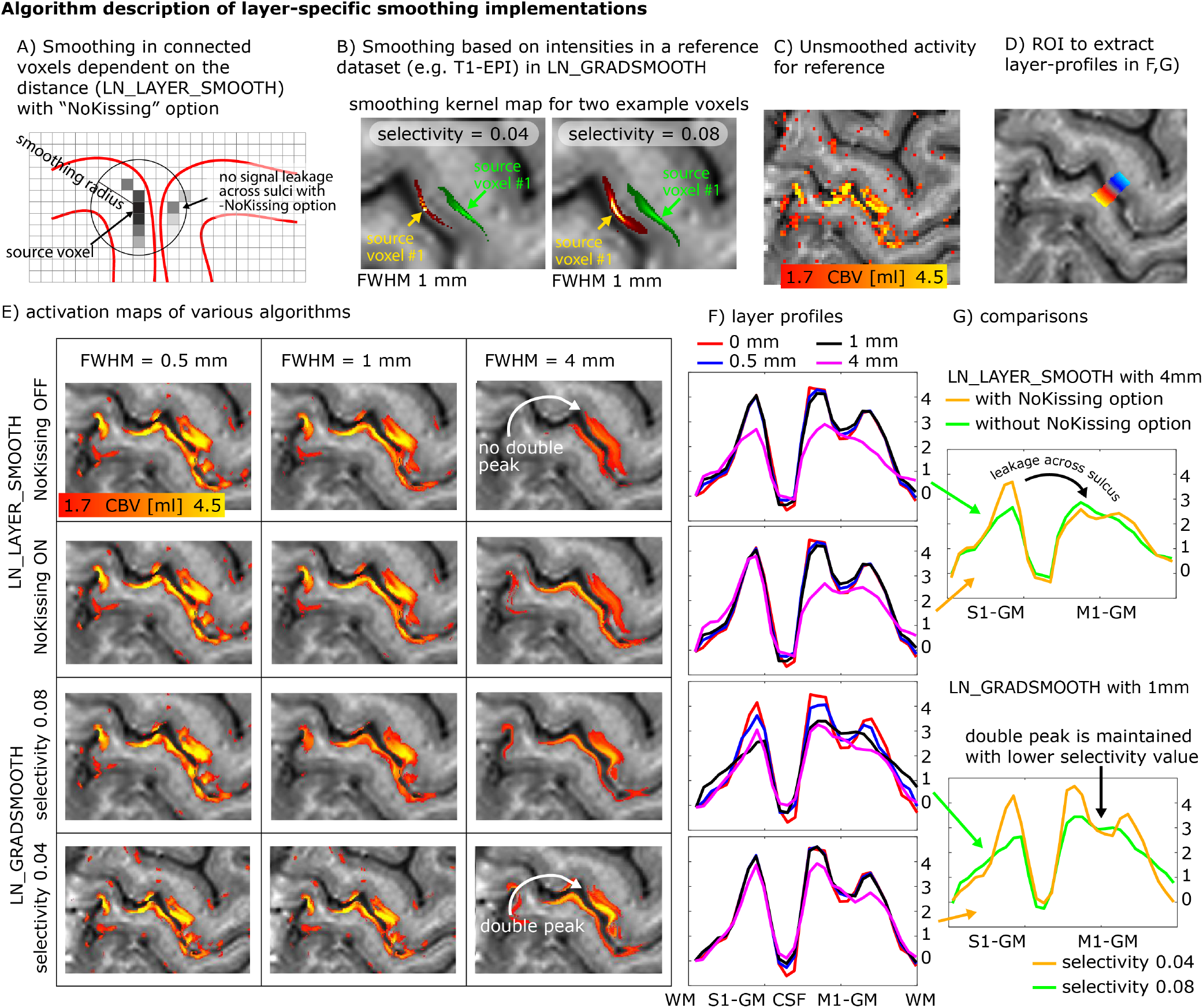
Layer-specific smoothing. Panels **A-B)** describe two different algorithms of layer-specific smoothing. Panel A) depicts that for spatial smoothing in the folded cortex, an Euclidean distance metric alone might not be a good enough estimate of columnar distances and cannot prohibit signal leakage across kissing gyri. Panel B) depicts example smoothing kernels in contrast-specific smoothing (anisotropic smoothing a.k.a. diffusion filter). Here the mean VASO EPI signal intensity with its inherent *T*_1_ -weighting is used as a reference to generate the voxel-wise smoothing kernel. The elongated shapes of the smoothing kernels depict the weighted signal leakage to voxels with similar anatomical contrast for two example voxels in green and red. The length of the kernel is determined by the FWHM parameter, while the width is determined by the *selectivity* parameter. Note that those Kernels are highly anisotropic. Panels **C-G)** exemplify the different results of the respective smoothing methods for an example dataset that comes with the software package. Here the characteristic double-stripe pattern is used as a layer-specific feature that is aimed to be preserved. Panel G) depicts that for extensive signal smoothing with FWHM of 4mm, layer-specific activation can leak across kissing gyri, if not explicitly prevented. Panel G) furthermore shows that the layer-specific double peak is only preserved for very conservative *selectivity* values. Note that the contrast specific smoothing algorithm with conservative *selectivity* values preserves the double peak layer activation pattern even for FWHM values of 4mm.

While this form of intracortical smoothing requires accurate knowledge of the local layer geometry within the cortex, there are alternative approaches to apply layer-specific smoothing in preferred layer-directions without the need to have pre-defined layer labels available. For example, it has been suggested to apply intracortical non-isotropic smoothing based on the local functional activation (Lohmann et al. 2018; Smith and Brady 1997) or based on the non-isotropic MRI signal intensity across the cortical depth. In LayNii, this form of smoothing is implemented in the program `LN_GRADSMOOTH` by taking advantage of the fact that high-resolution EPI intensities contain informative anatomical information too. This form of smoothing estimates the signal leakage kernel with two penalty functions; a) Gaussian weighted Euclidean distance of seed and target voxel, and b) local entropy weighted signal difference of the seed and target voxel. Just like for isotropic and layer-smoothing, the size of the Gaussian kernel can be adjusted by means of a user-defined `-FWHM` parameter in units of millimeter. The entropy weighted signal difference can be adjusted by the user by means of a `-selectivity` parameter. E.g., for a common `-selectivity` parameter of 0.08, the signal leakage is restricted to the local voxels that have a similar signal intensity, which differ from the central voxel by less than 8% of the local signal variance. For a more conservative `-selectivity` parameter of 0.04, the signal leakage is restricted to the local voxels that have a similar signal intensity, which differ from the central voxel by less than 4% of the local signal variance (see examples in Fig. 6B). The optimal selectivity value depends on the image’s contrast, the acceptable blurring across voxels with similar contrast, and the desired smoothness parameter (FWHM). Since the smoothing kernel is determined in a data-driven way, its application is significantly more convenient for the researcher to use. Labor intensive segmentation is not necessary to be able to perform this way of smoothing.

This algorithm of intensity-based smoothing is very similar to a large number of previously described edge-preserving smoothing methods (Gerig et al. 1992; Lohmann et al. 2018; Smith and Brady 1997; Weickert and Scharr 2002), which are implemented in AFNI, FSL, and CBS tools (which is also inherited by the Nighres wraper) as `3danisosmooth, LaminarIterativeSmoothing`, or `SUSAN`, and some of them are also referred to as *diffusion filter*. The LayNii implementation `LN_GRADSMOOTH` differs from those algorithms in the sense that it is optimized for applications in layer-fMRI. Since layer-fMRI data are usually very noisy, the smoothing kernel should not be generated on the actual time series data or the statistical activation data themselves. Instead, the LayNii implementation estimates the smoothing kernel on an independent optimized input file (given to `LN_GRADSMOOTH` with the `-gradfile` parameter), e.g., a mean EPI image with anatomical contrast (T1-EPI). This is readily doable in layer-fMRI data because of the exquisite structural details visible in sub-millimeter EPI. Furthermore, the LayNii implementation can avoid signal leakage of unconnected islands of similar signals intensities across kissing gyri (Fig. 6A).

This form of signal intensity based smoothing can also be used in combination with layer masks as described for `LN_LAYER_SMOOTH`. In this case, the originally hard borders of smoothing across layer-bins can be softened with larger `-selectivity` parameter values.

These forms of anatomically informed gradient smoothing and layer smoothing are particularly beneficial in layer-fMRI studies where it can be assumed that neighboring columns perform the same neural task with the identical layer-dependent activation signature (Beckett et al. 2020; Finn et al. 2019). And they are counter-productive, when signal differences of neighboring columnar structures are of interest.

Fig. 6C-G shows representative example results of anatomically informed smoothing applications. It can be seen that the characteristic double peak in the primary motor cortex is preserved across a wide range of algorithm parameters. Furthermore, it can be seen that for larger smoothing kernels, it becomes important to prevent signal leakage across kissing gyri.

Additional descriptions of the smoothing algorithms and their respective application in LayNii is given here: https://layerfmri.com/anatomically-informed-spatial-smoothing/.

### 2.5. Time series quality measures

Rigorous quality assessment of fMRI time series has become a fundamental pillar of any good code of conduct in fMRI research. In conventional fMRI, a consensus about good practices has been found and these practices are implemented in the major standardized analysis streams (Esteban et al. 2019). In layer-fMRI, however, conventional QA metrics are not sufficient or can even be completely inapplicable. Examples of metrics insufficient for layer-fMRI QA include a.) tSNR, b.) motion displacement, or c.) variance explained of neural ICA components.

- **tSNR:** Since tSNR is calculated by means and the temporal standard deviation (STDEV), it cannot easily capture multiple sources of variance that are adding non-linearly (sum of squares). Since layer-fMRI is usually dominated by thermal noise, layer-fMRI specific artifacts are not well captured in tSNR maps. They are rather hidden in the thermal noise.
- **Motion displacement:** While it is critically important in layer-fMRI to manually check displacement estimates like in conventional fMRI, it is not sufficient. In layer-fMRI motion usually comes along with higher-order EPI-Phase inconsistencies too. Namely, even when the rigid head motion can be corrected for, motion-related distortion changes and motion related B0-changes cause intermittent artifacts.
- **Variance explained in neural ICA components:** While the relative variance explained by neural ICA (independent component analysis) components is a useful measure for conventional fMRI, it is limited in layer-fMRI applications. ICA works best, if Gaussian and non-Gaussian fluctuation sources can be separated from each other. Unfortunately however, in layer-fMRI, the neurally-driven temporal fMRI signal fluctuations are deeply hidden in the dominating Gaussian thermal noise and it is not uncommon that there isn’t a single neural ICA component selectable in layer-fMRI analyses.

Due to the limits discussed above, conventional QA metrics in layer-fMRI data, additional and adapted metrics are needed. Since most layer-fMRI data are acquired in the thermal noise dominated regime, the noise can often be considered to be Gaussian. This means that non-thermal noise (e.g. artifact-induced signal clutter) can be identified by its non-Gaussian characteristics. Two useful measures to quantify how non-Gaussian a noise distribution is, are the measures of kurtosis and skewness (Berman et al. 2020). Fig. 7E,H depict a layer-fMRI example, where skew and kurtosis maps can depict artifact affected areas that are not clearly detectable in conventional tSNR maps.

**Figure 7.**
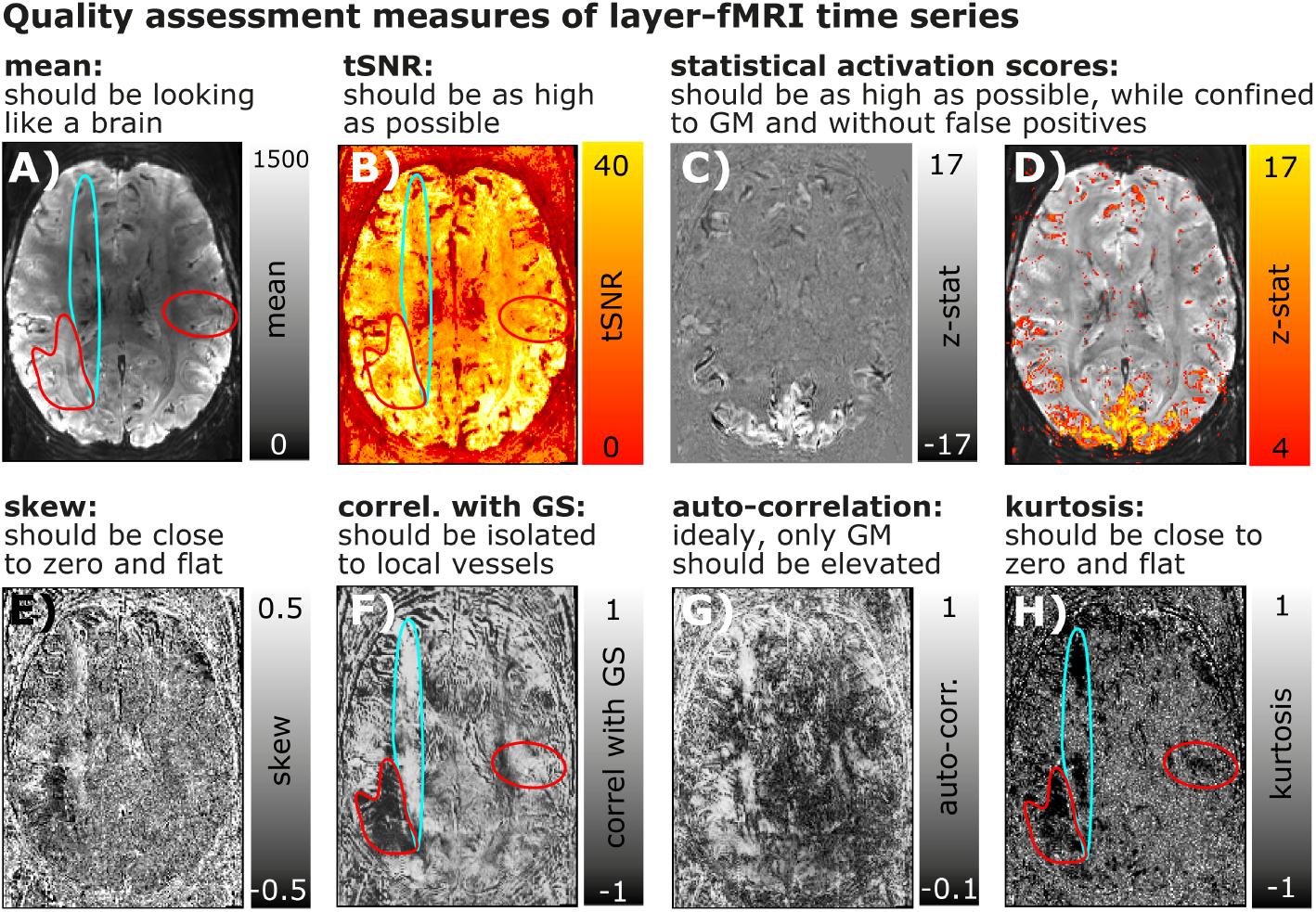
Conventional and layer-fMRI-specific quality metrics of time series data. Panels **A-D)** depict maps of conventional fMRI QA metrics of Mean, tSNR, and activation scores. Panels **E-H)** depict higher-order QA metrics that can be informative in layer-fMRI. While the QA measures in the top row suggest that the underlying time series is of high quality and that it is not severely limited by artifacts, higher-order QA measures in the bottom row show that there are indeed typical phase errors present in this example time series. For example, the turquoise and red marked areas highlight locations of non-Gaussian signal clutter that are likely referring to EPI phase problems during the readout. While these artifacts are clearly visible in most quality metrics of E)-H), they are hardly visible in first order quality metrics A)-D).

While thermal noise is not coupled across space and time, some forms of layer-fMRI artifacts result in strong noise coupling. This noise coupling can be used to identify artifacts and depict them despite the presence of dominating Gaussian noise. Informative measures that can be estimated on a voxel-wise level are: a) auto-correlation and b) the correlation with the global signal (GS). Fig. 7F,G depicts layer-fMRI examples where these measures can isolate and depict phase inconsistencies that are particularly limiting in layer-fMRI acquisition protocols.

While the above QA metrics are very useful to characterize the time course quality of individual voxels, they cannot easily capture spatio-temporal interactions. E.g. if neighboring voxels share the same sources of noise (e.g. due to spatial signal leakage), the above mentioned QA metrics are ignorant to this. This is particularly problematic because the EPI readout for layer-fMR is usually very long and layer fMRI data are, thus, expected to suffer from unwanted 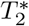-related temporal noise coupling between neighboring voxels.

To characterize the strengths and the pattern of the temporal correlation of neighboring voxels, a so-called noise kernel calculation is implemented in LayNii as described in Fig. 8A. The LayNii program estimates the average temporal correlation of neighboring voxels. In contrast to functional smoothness estimations in alternative software suites, the LayNii implementation of the noise kernel estimation does not simplify the noise coupling with individual Gaussian FWHM values and/or exponential decay terms. Instead, LayNii also writes out the 3-dimensional noise kernel as a volume nii file. This can be advantageous because a full noise kernel can capture negative side lobes, anti-correlations, as well as common saw-tooth patterns between odd and even lines. Furthermore, it also provides estimates along the diagonal directions, which can be helpful for 3D acquisition sequences (3D-EPI or 3D-GRASE) (Van Der Zwaag et al. 2012). This form of three dimensional noise-correlation kernel can in the future be used for data-driven deconvolution-based image sharpening filters (e.g. see the ‘-laurenzian’ flag in the LayNii program `LN_DIRECT_SMOOTH`). Furthermore, such analyses could be applied across layers and columns to estimate the topographical features of functional connectivity within the cortex (see Fig. S8 in (Huber et al. 2020b)).

**Figure 8.**
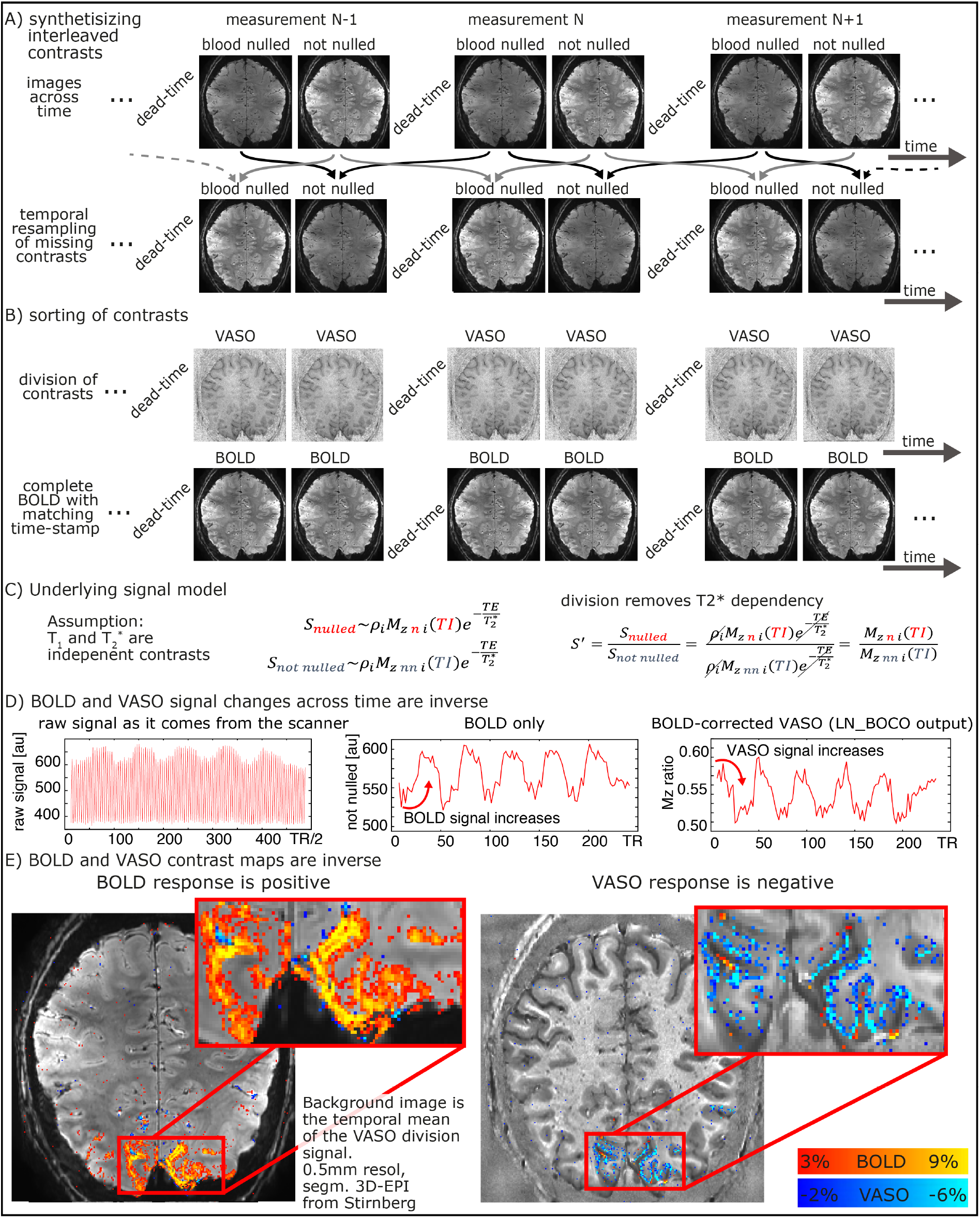
Spatiotemporal noise kernel. Panel **A)** depicts the major algorithm steps to estimate the noise kernel. Panel **B)** depicts representative results of the noise kernel in the whole-brain VASO layer-fMRI. It can be seen that the PSF in the second phase encoding direction has negative sidelobes, which suggests that the PSF is not well characterizable with FWHM estimates. The LayNii program `LN_NOISE_KERNEL` estimates the noise kernel for any time series dataset and writes it out as a three dimensional nifti file. These noise kernel files can then be used to characterize the quality of the time series. E.g., the user can then browse through several noise kernels from multiple layer-fMRI protocols and make judgements about which one is the best. For example, the user can ask questions about which protocol has the least 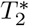 blurring (pink line in step 6 of panel A), and/or which flip angle scheme in 3D-EPI would result in the smallest negative point-spread function side lobes (blue dotted outline in panel B).

Additional descriptions of the QA algorithms, the limits of their interpretability and their respective application in LayNii is given here: https://layerfmri.com/qa/.

### 2.6. VASO specific programs

The VASO contrast (Lu et al. 2003) is believed to be inversely proportional to CBV changes and can non-invasively capture micro-vascular layer-specific signal responses through selective detection of signal changes in the extravascular volume compartment. The specific UHF-optimized Slab-selective slice inversion (SS-SI)-VASO approach acquires blood-nulled images and GE-BOLD images concomitantly (Huber et al. 2014b). To directly provide a CBV and BOLD contrast time course comparable to conventional fMRI signal contrast analyses, the raw VASO time series need to be temporally resorted and orthogonalized into clean 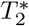-dependent and *T*_1_-dependent functional contrasts. The corresponding preprocessing steps are illustrated in Fig. 9A. To remove unwanted 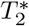-related signal in CBV-time series, LayNii assumes a simple magnetization model that obeys the Bloch equations (Fig. 9C). According to the Bloch equations, *T*_1_ and 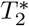 decays behave like two independent relaxation effects that can be described in a multiplicative fashion. This means that unwanted 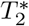 contrast in the CBV-weighted signal can be canceled out by means of dynamic division of MR images with and without preceding *T*_1_-weighting (Fig. 9C).

**Figure 9.**
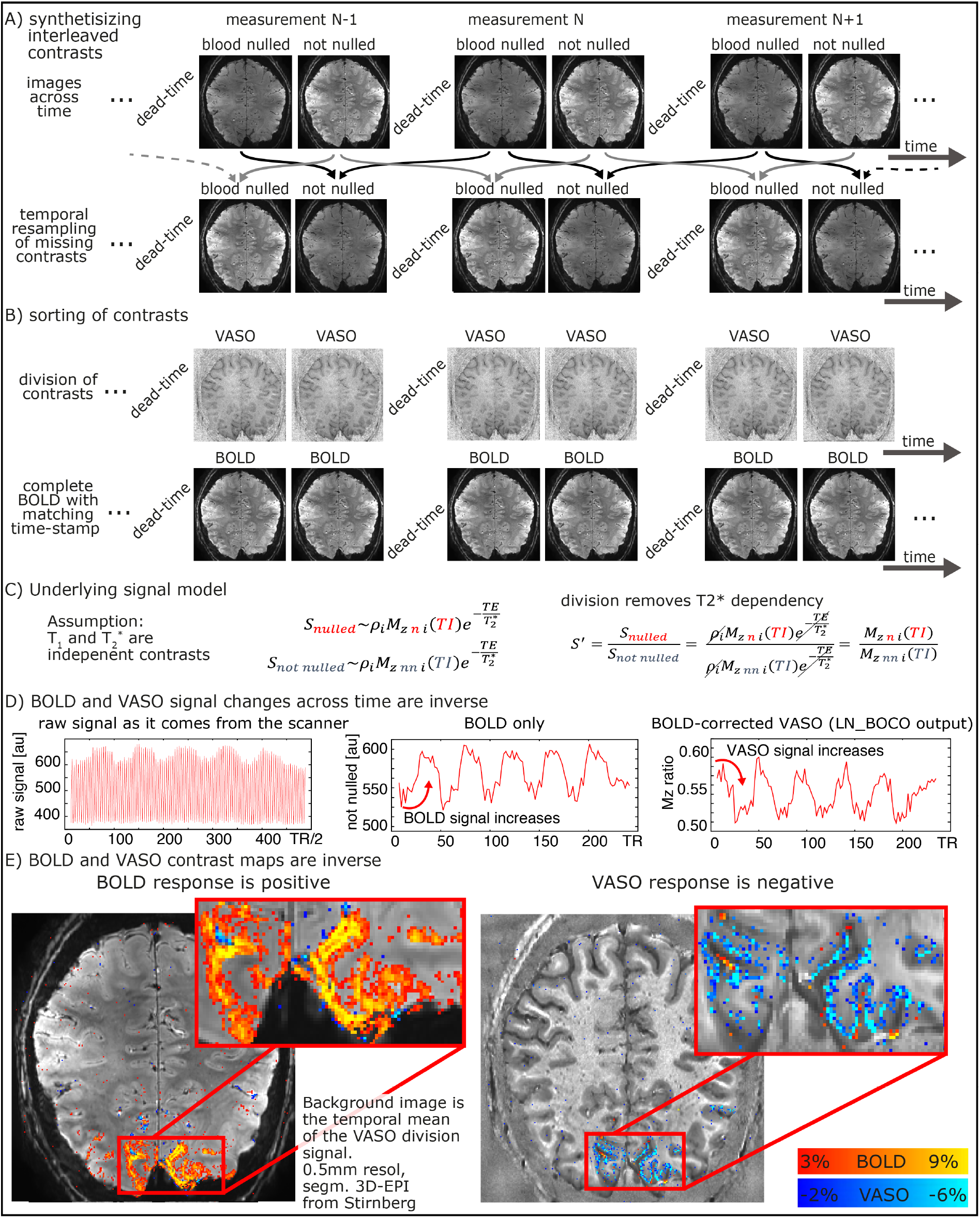
VASO processing at high magnetic fields with the LayNii program LN_BOCO. Due to the short 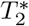 at UHF and the long readout of high-resolution EPI, all VASO data are contaminated with unwanted BOLD contrast. In layer-fMRI application of VASO, this is accounted for by concomitant acquisition of BOLD and VASO in the SS-SI-VASO approach, analogously to ASL-fMRI. The purpose of this figure is to illustrate the relevant processing steps in the LayNii program `LN_BOCO` to correct for unwanted BOLD contamination in VASO. Panels A-B) depict how the original time series of alternating images with and without blood nulling are temporally interpolated and sorted into two parallel time series of respective contrasts only. Panel C) depicts that in SS-SI-VASO, the BOLD contamination is believed to be a multiplicative factor and be taken care of with a division operation. Panels D-E) depict a representative functional dataset of a flickering checkerboard experiment. It can be seen in the time courses that VASO is anti-correlated to BOLD. While the BOLD signal shows a signal increase during activation, upon BOLD correction, VASO shows a signal decrease. The data presented here have been acquired on a SIEMENS Terra in Glasgow with a segmented-EPI sequence from Ruediger Stirnberg (**?**).

Fig. 9D-E depicts a representative example of VASO and BOLD data as they are generated from LayNii. Note that the VASO output is inversely correlated with dynamic CBV changes.

For a hands-on example of how the application of LayNii works for layer-fMRI VASO applications see here: https://layerfmri.com/analysispipeline/.

### 2.7. Model based deveining

Even though high-resolution GE-BOLD is known to suffer from unwanted locally-nonspecific signals of large draining veins and the location of large draining veins on the cortical surface, due to its high sensitivity it is the most popular acquisition method for layer-fMRI data. Due to the blood drainage towards the cortical surface (Olman et al. 2007; Petridou and Siero 2019), the unwanted GE-BOLD signal of large veins is known to increase towards the cortical surface. Based on the angioarchitecture, it has been proposed that the unwanted layer-specific venous signal bias in GE-BOLD can be accounted for in retrospective model-based correction. A number of models are popular in the field (Fig. 10A):

**Figure 10.**
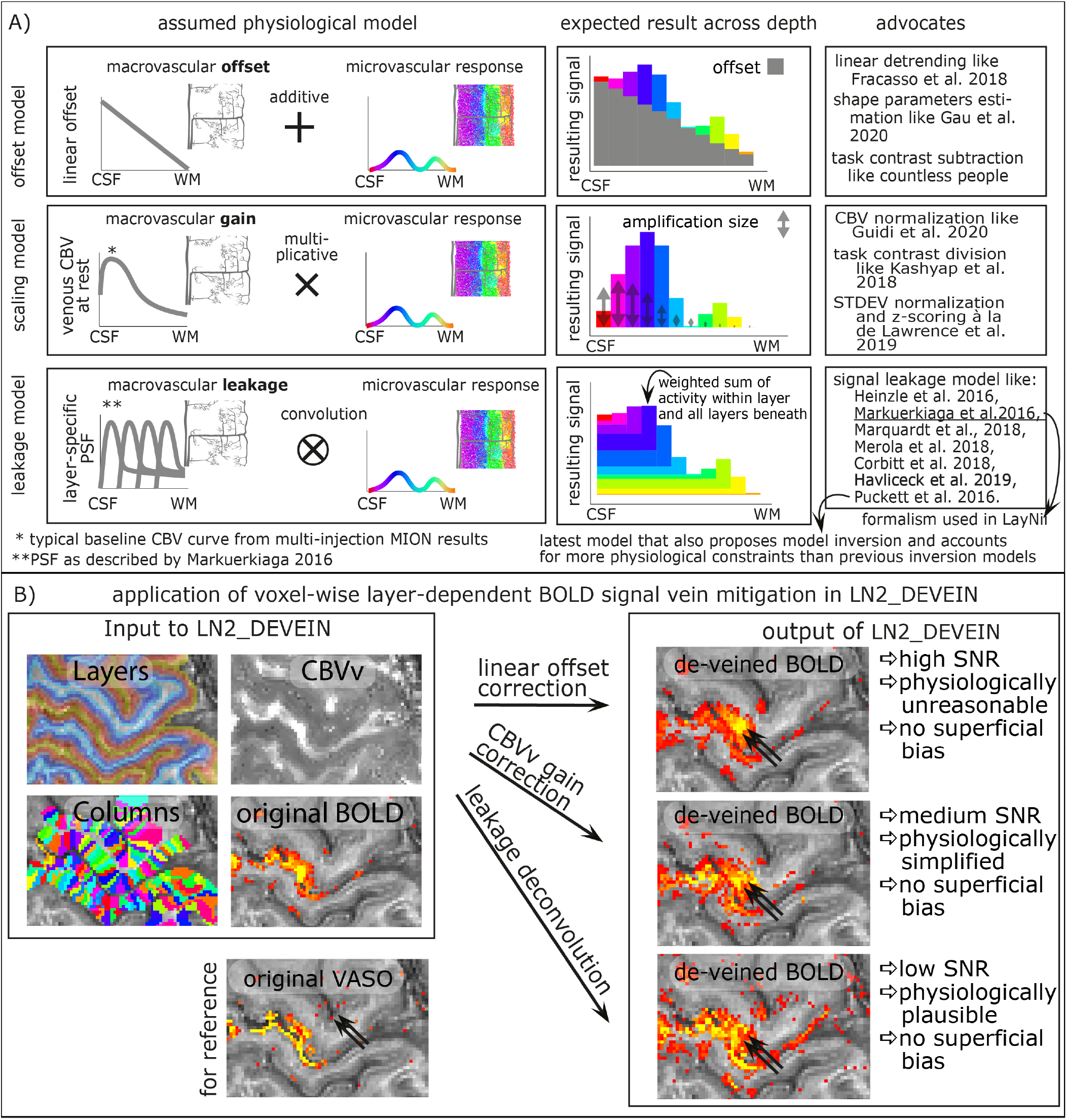
Layer-dependent model-based deveining strategies. The three most often used strategies of layer-dependent vein mitigation are based on a linear offset model, a scaling model, or a leakage model. The respective models are illustrated in panel A. While all models can be used to predict the increasing GE-BOLD signal towards the cortical surface, their assumed physiological signal origin and the corresponding vein mitigation algorithm is fundamentally different. Panel B exemplifies the application of layer-dependent vein mitigation in LayNii and depicts representative results.

- The **linear-offset model** (Fracasso et al. 2018; Gau et al. 2020) is based on the assumption that the layer-specific microvascular signal is added on top of a task-independent macrovascular signal. Thus, it is assumed that a simple task-contrast subtraction or a simple linear-trend-removal would get rid of the unwanted macrovascular component of the BOLD signal. The linear offset model has been explicitly discussed in the context of linear correlation analyses (Fracasso et al. 2018) and in the context of layer-dependent shape parameters (Gau et al. 2020). This linear model is furthermore implicitly used for in countless task contrast subtraction analyses of previous layer-fMRI studies.
- The **scaling model** (Guidi et al. 2020; Kashyap et al. 2018; Kazan et al. 2017; Lawrence et al. 2019) is based on the assumption that the layer-specific bias in GE-BOLD is driven by layer-dependent variations of vein density. As such, it is assumed that the superficial signal is larger than the signal in deeper layers, simply because the amount of venous blood volume is higher in superficial layers compared to deeper layers. Or in other words, the macrovasculature acts like a layer-specific signal amplification (gain). Thus, it is assumed that a simple multiplicative (or divisory) normalization can correct for the macrovascular signal bias. This layer-dependent signal normalization has been proposed as part of several layer-fMRI analysis strategies, including:

a. scaling the layer-dependent signal with estimates of layer-dependent venous CBV (Guidi et al. 2020; Kazan et al. 2017; 2016),
b. normalizing the signal difference between different task responses by the mean signal response of all involved task responses (Kashyap et al. 2018),
c. refraining to infer neuroscience conclusions solely based on task response differences in favor of focusing on task response ratios that are normalized by the presumably vascularly driven signal fluctuations (Lawrence et al. 2019).
- The **leakage model** (Markuerkiaga et al. 2016) is based on the assumption that the BOLD signal in each layer constitutes a signal mixture of activity originating in multiple layers. Namely, it is assumed that a voxel in a given layer contains some signal from the layer of interest plus an additional unwanted integrated signal of all the layers beneath. E.g. it is assumed that the BOLD signal in a voxel of a superficial layer contains both the desired superficial signal plus unwanted signal from the deeper layers. The BOLD signal model in each layer is then usually parameterized as a weighted sum of the layer itself and the signal from deeper layers. To use this model to correct for unwanted leaked signals, a spatial signal deconvolution approach is applied. (Note that some implementations of such deconvolution models refrain from the term “deconvolution” in favor of the terms “matrix inversion” or “consecutive signal subtraction”). Very often, the leakage model is combined with the scaling model (Havlicek and Uludag 2019; Markuerkiaga et al. 2016; Merola and Weiskopf 2018). Layer-fMRI focused implementations of BOLD signal models that assume spatial signal leakage across layers have been described by a large number of research labs, including models from Heinzle et al. 2016, Markuerkiaga et al. 2016, Merola et al. 2018, Puckett et al. 2016, Lacy et al. 2020 Corbitt et al. 2018, and Havlicek et al 2019. All of these signal leakage models estimate the signal in superficial layers as a sum of the microvascular response within the given layer plus a weighted sum of the signal from all other layers below. The main difference between the various leakage models comes from the procedure by which the respective summation weights are estimated. The model in (Lacy et al. 2020; Puckett et al. 2016) describes the venous signal draining with travelling wave equations. The models described in (Corbitt et al. 2018; Markuerkiaga et al. 2016) perform forward simulations without the explicit aim to invert the model for venous signal removal. The models described in (Havlicek and Uludag 2019; Heinzle et al. 2016), propose such a model inversion for the removal of unwanted venous signal in layer-fMRI signal processing. The newer model in (Havlicek and Uludag 2019) entails more physiologically-informed constraints than the model in (Heinzle et al. 2016). In most of those models, the summation weights of layer-dependent signal leakage are derived based on more-or-less appropriate assumptions of the layer-dependent blood vessel architecture and/or experimental data. The weights usually depend on resting CBVv and CBF assumptions across cortical depth.

In current layer-dependent applications that use model-based deveining, the models are applied area-wide (e.g. (Marquardt et al. 2018)). I.e., fMRI signals are pooled from the layers of large cortical patches and their effect is solely investigated in the form of one-dimensional layer profiles. In LayNii, however, the various vein mitigation algorithms work on a voxel-by-voxel level. This allows the researcher to appreciate the signal distribution along the topographical space across layers and columns, simultaneously. Furthermore, the resulting signal maps provide an intuitive understanding of the noise amplification that comes along with the various deveining algorithms of the respective models (See Fig. 10B for examples).

All three model-based deveining methods are included in LayNii. The linear offset model is used by executing `LN2_DEVEIN` with the flag `-linear`. The CBV scaling model is used by executing `LN2_DEVEIN` with the flag `-CBV`. The leakage model is used if `LN2_DEVEIN` is executed without either of those flags. The formalism of implementation of the leakage model is largely based on Markuerkiaga et al. 2016. Despite the different approach to estimate leakage weights, the formalism implemented in LayNii that describes the signal leakage as a spatial weighted signal summation is identical for all published leakage models mentioned above. Since CBVv can be obtained from the GLM residuals (Guidi et al. 2020; Kazan et al. 2017; 2016), no additional measurements are needed. Additional descriptions of the model-based deveining algorithms and their respective application in LayNii is given here: https://layerfmri.com/devein/. Note that the model-based deveining described here should not be confused with the auxiliary method of (semi-)automatic detection of large veins (Bause et al. 2020) and masking of vein-dominated voxels (Moerel et al. 2018a).

#### 2.7.1. Limits of model based deveining

While the authors are excited about the emergence of model-based deconvolution methods in many research labs, these model-based vein mitigation methods are just starting to be validated, their shortcomings are just starting to be understood, and their applications are not widely established except for exploratory use. Currently, it is still more established in the field to avoid unwanted venous signals by means of advanced acquisition strategies, rather than adding additional signal analysis (deconvolution) steps. Specific extensions to current models that are currently being investigated are:

- So-called blooming effects (cross-voxel dipole fields) of very large draining veins into neighboring voxels and layers that are currently not accounted for in model-based deveining methods (Bause et al. 2020; Moerel et al. 2018a).
- More detailed knowledge of the signal distribution between intra and extravascular BOLD effect across field strengths and echo times for various readout trajectories during 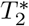-decay needs to be obtained, validated, and incorporated in future models. Due to the sheer numbers of model parameters and experimental setups, elaborate and comprehensive validation studies are needed.
- Orientation effects of pial and ascending veins with respect to the external magnetic field (Viessmann et al. 2019) that can lead to overestimation or underestimation of the contribution in model-based deveinig, if not accounted for.
- A crucial requirement of model-based deveining is the assumption of a good baseline signal. Thus, impractical inter-stimulus-intervals and task-duration dependencies might hamper straightforward application of model-based deveining in neuroscience studies.

## 3. 3. Discussion

Here, we present an fMRI analysis toolbox that is specifically developed for layer-fMRI applications. It has minimal requirements on the input data and works in the native data format of voxels as they come off the scanner. Its application ranges from layer/column assignment, over anatomically informed smoothing and QA, up to model-based deveining and VASO analyses.

### 3.1. What LayNii does not contain

LayNii is not designed to duplicate analysis features that have already been implemented in alternative fMRI software packages with the same capabilities. Even though there are specific layer-fMRI demands for partial coverage image alignment (Weldon et al. 2019; Weldon and Olman 2020), LayNii does not perform it itself. Since LayNii also works in the native EPI-space, image alignment is not absolutely necessary to begin with. If a layer-fMRI user is interested in alignment constraints that are specific to layer-fMRI data, we refer to the instructions of how to perform image alignment of layer-fMRI data with ITK-snap (Yushkevich et al. 2006) and ANTs (https://layerfmri.com/high-quality-registration/), or with AFNI (https://blog.cogneurostats.com/aligning-partial-volume-fmri-in-afni/) and (Navarro et al. 2020). Furthermore, even though tissue type segmentation is particularly important in layer-fMRI analyses, LayNii does not contain its own segmentation tools. In layer-fMRI, GM borders often need to be manually adjusted and corrected anyway. Even though this is a time consuming and tedious task of several work days, in layer-fMRI, it is often most efficient to fully manually segment the areas of interest within the limited EPI coverage without the necessity of automated segmentation tools. For large coverages and/or a rough first estimate of GM borders as an input to LayNii, alternative software packages can be used. E.g. the interested reader is referred to an instruction of how to use FreeSurfer, nighres and SUMA to generate an input rim file for LayNii here: (https://layerfmri.com/getting-layers-in-epi-space/), this segmentation can be further corrected with the semi-manual segmentation tool Segmentator (Gulban et al. 2018) (https://github.com/ofgulban/segmentator). The LayNii program `LN_RIMIFY` can be used to convert the segmentation output of common third-party software tools to a LayNii-readable rim file.

A collection of non-LayNii scripts and programs that are designed to work in tandem with LayNii available to download here: https://github.com/ofgulban/LayNii_extras.

While it is a dedicated aim of the developers to include it in the LayNii software suite, the current version of LayNii does not contain a program that estimates the geometric angle between the cortical surface and the voxel grid or the main magnetic field.

### 3.2. Potential application of LayNii outside of layer-fMRI

Here, ‘layer-fMRI’ is used as an umbrella term for depth-dependent fMRI, intra-cortical fMRI, and sub-millimeter fMRI in general1. This means that LayNii is also explicitly intended for signal analyses of columnar structures (e.g. see Fig. 6)). LayNii is originally intended for - but not limited to-functional MRI. In fact, LayNii is applicable and has been applied to high-resolution structural MRI, as well as histology data with and without restricted field of views (E.g. Fig. 3 and (Huber et al. 2017)). We believe that LayNii might also be specifically suited for in-vivo sub-millimeter diffusion-weighted data. While LayNii is time-efficiently applicable to conventional high-resolution whole brain anatomical data, it is not specifically advertised for this purpose. While LayNii can be applied on conventional structural data with distortion-corrected and aligned functional data (Finn et al. 2019; Weldon and Olman 2020; Zaretskaya et al. 2020), there are alternative analysis software packages that are more extensively tested, documented and supported with exhaustive educational tutorials for this purpose.

While LayNii is specifically designed for high-resolution applications, individual programs can be applied to lower spatial resolution fMRI data too. Namely, the VASO-specific analyses can -and have been-applied to lower-resolution 3 mm data too (Huber et al. 2014b). Similarly, the quality assessment metrics can also be beneficial at lower resolutions (Kurban et al. 2020).

Since LayNii works directly in voxel space without imposing topological constraints, it can straightforwardly generate layers and column estimates in 2D data. This allows applications in common single slice data of preclinical MRI in rodents, cats and monkeys, and it allows layer profile extraction of figures from any electronic publication including the seminal microscopy images of an entire century of cyto-architecture (Brodmann 1909), myelo-architecture (Vogt and Vogt 1919) and angioarchitecture (Pfeifer 1940) research. Note, however, that in 2D, the equi-volume layering approach converges with an equi-area approach. For more information on how to generate layer-profiles of literature figures see: https://layerfmri.com/how-to-convert-any-paper-figure-into-a-layer-profile/.

### 3.3. Usage in the field

The LayNii software suite has been used in multiple high-resolution fMRI studies across the world. It’s Github site welcomes >1400 unique visitors per year and its code is being cloned >100 times per month. It found particular application for layer-fMRI studies in the motor cortex (Huber et al. 2017), in sensory cortex (Yu et al. 2019), DLPFC (Finn et al. 2019), across association cortices (**?**) for columnar imaging in the motor cortex (Huber et al. 2020b), in layer-specific functional connectivity mapping (Huber et al. 2020a), for mental imaginary layer-fMRI (Persichetti et al. 2020), for methods development of new sequences (Beckett et al. 2020; Chai et al. 2019; Guidi et al. 2020), for visual layer-fMRI (Zamboni et al. 2020), for model-based removal of vein effects in layer-fMRI-EEG (Marsh et al. 2020) and for methods debugging of human 9.4T layer-fMRI (Huber et al. 2018). In the early days of LayNii, its programs were shaped and optimized by continuous interactions with its users. Current and future requests for new features and support are encouraged via LayNiis github issue page.

### 3.4. How LayNii evolved from other packages

All the discussed LayNii programs (BOLD correction, equidistance and equi-volume layering, etc.) have been initially implemented in the ODIN (Object-oriented Development Interface for NMR) environment (Jochimsen and Von Mengershausen 2004). This LayNii predecessor software suite was originally used in many initial layer-fMRI studies (Guidi et al. 2016; 2020; Huber et al. 2014a; 2017; 2016a;b). However, since this software package had more than 30 dependencies, and required files to be located in certain root-locations, it was not straightforward to use across various operating platforms, nor was it installable on servers with conservative user access. Thus, with guidance from the NIH data science and sharing team (https://cmn.nimh.nih.gov/) and the AFNI team (https://afni.nimh.nih.gov/), the layer-fMRI algorithms were reimplemented outside of ODIN with a data I/O that is inherited from the original NIFTI release (https://nifti.nimh.nih.gov/). Thus, LayNii does not have any residual external dependencies on third-party software, nor does it rely on libraries.

## Conclusion

Here we introduce a new dedicated software toolbox for layer-specific (functional) MRI. It is open source and available for installation via source code or pre-compiled binaries for Linux, Windows and macOS and provides a comprehensive set of advanced techniques for layer-fMRI analyses. The current functionality largely focuses on applications where the challenges of layer-fMRI data do not allow the application of standard analysis pipelines of the major software packages. Though, we hope that the modular package structure facilitated augmentation of analysis pipelines that also contain other software packages.

## Acknowledgements

We thank Pilou Bazin for contributing comments, discussions and ideas regarding the algorithms and implementation of the equi-volume layerification and layer-specific smoothing programs of LayNii. We thank Kamil Uludag and Martin Havlicek for comments on the manuscript regarding the model-based vein mitigation.

## Funding

Parts of this research was supported by the NIMH Intramural Research Program (ZIA-MH002783). Laurentius Huber was funded form the NWO VENI project 016.Veni.198.032 for part of the study. Benedikt Poser is partially funded by the NWO VIDI grant 16.Vidi.178.052 and by the National Institute for Health grant (R01MH/111444) (PI David Feinberg). Portions of this study used the high-performance computational capabilities of the Biowulf Linux cluster at the National Institutes of Health, Bethesda, MD (biowulf.nih.gov). Rainer Goebel is partly funded by the European Research Council Grant ERC-2010-AdG 269853 and Human Brain Project Grant FP7-ICT-2013-FET-F/604102. Nils Nothnagel and Jozien Goense are funded by the Medical Research Council (MR/R005745/1). Andrew Tyler Morgan is funded by the Medical Research Council (MR/N008537/1) and the European Union’s Horizon 2020 Framework Programme for Research and Innovation under the Specific Grant Agreement No. 785907 and 945539 (Human Brain Project SGA2 and SGA3)

## Discussions of code

We thank Sriranga Kashyap for many informative questions of previous versions of LayNii. We thank Kamil Uludag for helpful discussions and contributions on model-based deveining (Fig. 10). We thank Elisha Merriam for discussions about the least noise amplifications in `LN_BOCO` with spline surround-division. We thank the many users of LayNii that submitted bug reports and feature requests to our repository.

## Acquisition of example data

We thank Kenny Chung and Joe Stolinski for radiographic assistance with experiments conducted at NIH (Figs. 4C, 5E, 6, 9B). We thank Sean Marrett for co-acquiring data used in Fig. 6, and 9B. We thank FMRIF (especially Andy Derbyshire), NMRF (especially Joelle Sarlls and Lalith Talagala), and Scannexus (especially Chris Wiggins), where the example data were acquired. We thank Ruediger Stirnberg and Tony Stoeker for their segmented IR-3D-EPI sequence that was used to obtain data in Fig. 8.

## Financial interest

Author OFG and RG work for Brain Innovation and have financial interest tied to the company.

## Beta users

We thank the many LayNii users who sent feedback via Github issues. We also thank Dimo Ivanov, Sri Kashyap and Deni Kurban for testing the binaries.

## Data and Software availability

Example data, source code, binary executables, installation instructions, installer packages can be found here: https://doi.org/10.5281/zenodo.3514298 via a BSD-3 license. A Dockercontainer with preinstalled LayNii and example data is avaliable here: https://hub.docker.com/r/layerfmri/laynii_v2.0.0. Also see common analysis pipelines in which LayNii is used here: https://github.com/ofgulban/LayNii_extras.

## Footnotes

1 https://layerfmri.com/terminology/

## Notes

https://doi.org/10.5281/zenodo.3514298

https://github.com/layerfMRI/LAYNII/tree/master/test_data

### Summary of Updates

The major revision contains: -> new section is included about the limits of model-based devenining -> rewriting of Conclusion section -> additional data are shown for layerification on different spatial grids. -> it is emphasized what the software suite does not contain.

